# A neural crest stem cell-like state drives nongenetic resistance to targeted therapy in melanoma

**DOI:** 10.1101/2020.12.15.422929

**Authors:** Oskar Marin-Bejar, Aljosja Rogiers, Michael Dewaele, Julia Femel, Panagiotis Karras, Joanna Pozniak, Greet Bervoets, Nina Van Raemdonck, Dennis Pedri, Toon Swings, Jonas Demeulemeester, Sara Vander Borght, Francesca Bosisio, Joost J. van den Oord, Isabelle Vanden Bempt, Diether Lambrechts, Thierry Voet, Oliver Bechter, Helen Rizos, Mitch Levesque, Eleonora Leucci, Amanda W. Lund, Florian Rambow, Jean-Christophe Marine

**Affiliations:** Laboratory for Molecular Cancer Biology, Center for Cancer Biology, VIB, Leuven, Belgium; Laboratory for Molecular Cancer Biology, Department of Oncology, KULeuven, Leuven, Belgium; Department of Cell, Developmental & Cancer Biology, Knight Cancer Institute, Oregon Health and Science University, Portland (OR); VIB Technology Watch, Technology Innovation Lab, VIB, Leuven, Belgium; Laboratory of reproductive genomics, Department of Human Genetics, KU Leuven, Leuven, Belgium; Department of Pathology, UZLeuven, Belgium; Laboratory of Translational Cell and Tissue Research, Department of Pathology, KULeuven and UZ Leuven, Leuven, Belgium; Center for Human Genetics, KULeuven, Belgium; Laboratory of Translational Genetics, Center for Cancer Biology, VIB, Leuven, Belgium; Laboratory of Translational Genetics, Center for Human Genetics, KULeuven, Belgium; Department of General Medical Oncology UZ Leuven, Belgium; Macquarie University, Sydney, NSW, Australia; Melanoma Institute Australia, Sydney, NSW, Australia; Department of Dermatology, University of Zürich Hospital, Zürich, Switzerland; Laboratory for RNA Cancer Biology, Department of Oncology, LKI, KU Leuven, Leuven, Belgium; Trace PDX platform, Department of Oncology, LKI, KU Leuven, Leuven, Belgium

**Author notes:** Correspondence should be addressed to Florian Rambow or Jean-Christophe Marine. Lead contact: Jean-Christophe Marine.

**Keywords:** therapy resistance, nongenetic reprogramming, cutaneous melanoma, Neural Crest Stem Cells, FAK/SRC signaling

## Abstract

The ability to predict the future behaviour of an individual cancer is crucial for precision cancer medicine and, in particular, for the development of strategies that prevent acquisition of resistance to anti-cancer drugs. Therapy resistance, which often develops from a heterogeneous pool of drug-tolerant cells known as minimal residual disease (MRD), is thought to mainly occur through acquisition of genetic alterations. Increasing evidence, however, indicates that drug resistance might also be acquired though nongenetic mechanisms. A key emerging question is therefore whether specific molecular and/or cellular features of the MRD ecosystem determine which of these two distinct resistance trajectories will eventually prevail. We show herein that, in melanoma exposed to MAPK-therapeutics, the presence of a neural crest stem cell (NCSC) subpopulation in MRD concurred with the rapid development of resistance through nongenetic mechanisms. Emergence of this drug-tolerant population in MRD relies on a GDNF-dependent autocrine and paracrine signalling cascade, which activates the AKT survival pathway in a Focal-adhesion kinase-(FAK) dependent manner. Ablation of this subpopulation through inhibition of FAK/SRC-signalling delayed relapse in patient-derived tumour xenografts. Strikingly, all tumours that eventually escaped this treatment exhibited resistance-conferring genetic alterations and increased sensitivity to ERK-inhibition. These findings firmly establish that nongenetic reprogramming events contribute to therapy resistance in melanoma and identify a clinically-compatible approach that abrogates such a trajectory. Importantly, these data demonstrate that the cellular composition of MRD deterministically imposes distinct drug resistance evolutionary paths and highlight key principles that may permit more effective pre-emptive therapeutic interventions.

## Introduction

The inability to fully eradicate metastasis, the major source of cancer-related deaths, is one of the most important challenges faced by modern oncologists. The past two decades have brought a bevy of therapeutics that block immune checkpoints or interfere with cancer signalling pathways^1^. These agents have revolutionized patient care. Many patients with advanced metastatic disease now achieve objective responses. Unfortunately, however, the vast majority of patients who initially respond to treatment, later develop resistance. This is because all available therapeutic modalities almost invariably leave a reservoir of residual cancer cells behind, traditionally called Minimal Residual Disease (MRD), from which relapse inevitably emerges.

The most commonly accepted explanation for the inexorable evolution of resistance invokes genetic alterations and selection of mutant cells carrying a relevant mutation acquired by chance before or during treatment^2^. The large repertoire of mutation-driven evasion mechanisms observed in response to a given therapeutic challenge has highlighted the enormous plasticity of the cancer genome and raised scepticism about the actual effectiveness of precision oncology^3^. Devising therapeutic strategies that target all these resistance-conferring genetic events was compared to a whack-a-mole game, which is unwinnable considering that the vast majority of oncogenic mutations are not even druggable^4^. These data highlighted the need to improve effectiveness of treatment before mutational acquired resistance prevails.

Recent findings indicated that drug-tolerant phenotype(s) (as defined as the ability to survive the drug treatment) can be transiently acquired through non-mutational mechanisms^5-10^. Studying the response of BRAF-mutant melanoma to MAPK-targeted therapy, we recently reported the co-emergence of varying combinations of distinct drug-tolerant transcriptional states^6^. Four distinct melanoma drug tolerant states were identified: the Starved Melanoma Cell (SMC) state sharing transcriptomic features of nutrient-deprived cells^11^, a Neural Crest Stem-like Cell (NCSC) state, an invasive or mesenchymal-like state that was recently renamed undifferentiated state^12,13^ and a hyperdifferentiated state. Interestingly, the NCSC transcriptional program appeared to be largely driven by the nuclear receptor RXR and, consistently, an RXR antagonist (HX531) mitigated, but did not prevent, accumulation of NCSCs in MRD and delayed the development of resistance^6^. These data illustrated the potential of MRD-directed therapies^14^ and indicated that the pool of NCSCs can serve as the cellular origin of drug resistance. However, since no RXR antagonist has been approved by the FDA to date, translating this work into a new treatment in the short-term requires the identification of more efficient and clinically-compatible approaches to target these cells.

Moreover, although firm evidence for this is still lacking, an increasing body of data has indicated that drug resistance (which, as opposed to tolerance, designates the ability to proliferate despite therapy exposure) may also develop in absence of detectable genetic alterations in several cancers^15-22^. Genomic analyses have failed to identify a genetic cause for the development of stable MAPK-resistant melanoma cell cultures and lesions that regrew in drug-exposed PDXs and human patients^15,17,20,23,24^. Given that melanoma is the tumour type with the highest mutation load, these observations raise the possibility that nongenetic resistance may be a rather common phenomenon. An improved understanding of the mechanisms underlying nongenetic drug resistance may therefore yield impactful therapeutic strategies across tumour types^25-27^.

Importantly, whether therapy resistance can emerge from NCSCs (or any other drug tolerant cells) through genetic and/or nongenetic mechanisms remains unclear. A related, but more fundamental, question is whether the selection between these two distinct resistance trajectories occurs in a stochastic or deterministic manner.

## Results

To assess the extent to which nongenetic mechanisms contribute to the development of resistance to targeted therapy in melanoma we performed an *in silico* multicentric meta-analysis of whole-exome sequencing (WES) datasets from clinical samples (n=59) that progressed on combinations of BRAF- and MEK-inhibitors, one of the standards of care for BRAF-mutant melanoma patients (Figure 1A and TableS1). About 20% of the samples exposed to this treatment did not exhibit any evidence of single-nucleotide alterations (SNAs) previously validated as drivers of resistance to MAPK-therapeutics^23,24,28,29^ nor amplification of the *BRAF* gene, one of the most common resistance-conferring genetic alterations^30^.

**Figure 1:**
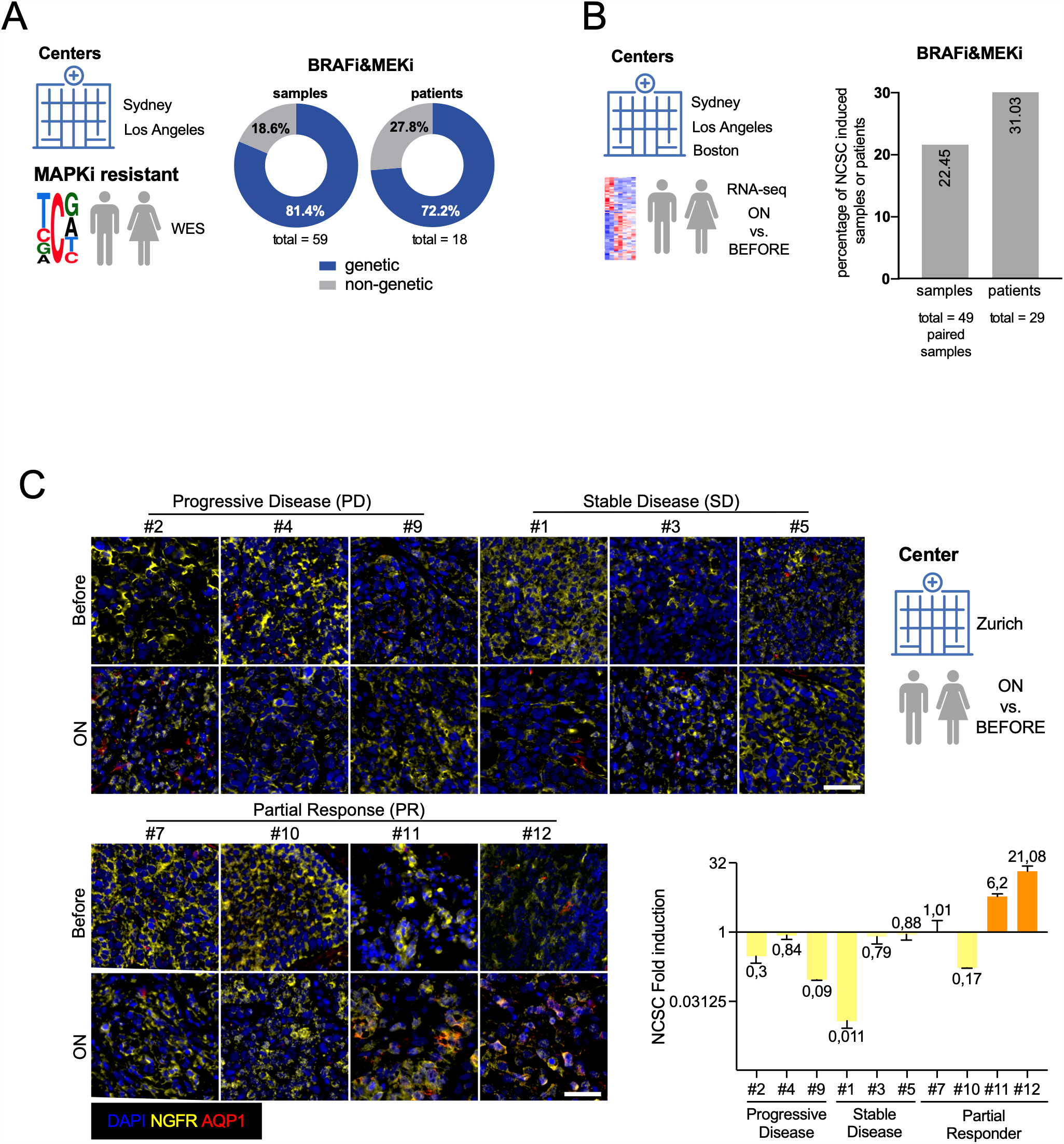
Non-genetic resistance to MAPK inhibition and the induction of the NCSC state are frequent events in human melanoma. A) MAPKi-resistant human melanoma samples were analysed by whole-exome sequencing (WES) for absence (non-genetic) or presence of genetic events^17,23,28^. The analysis included samples from the Melanoma Institute Australia (Sydney, Australia) and UCLA Dermatology (Los Angeles, USA). The percentages of samples or patients were determined for genetic and non-genetic resistance. B) Gene expression data (bulk RNAseq) of melanoma from drug naïve samples/patients and following exposure to MAPK-inhibitors were assessed for the induction of the neural crest stem like (NCSC) transcriptional program. The analysis included samples from the Melanoma Institute Australia (Sydney, Australia), UCLA Dermatology (Los Angeles, USA) and Massachusetts General Hospital (Boston, USA)^17,23,28,65^. C) Immunostainings show emergence of AQP1+/NGFR+ double-positive NCSCs upon MAPK-inhibition (O/T = ON treatment) in clinical samples from responders, but not from patients that progressed or exhibited stable disease ON treatment. Representative images are shown (the error bars represent standard deviations of counting n= 5 fields per tumour sample). The analysis included samples from University of Zurich Hospital (Zurich, Switzerland).

We previously identified the drug-tolerant NCSC population as a putative driver of melanoma recurrence^6^. Interestingly, *in silico* analysis of RNA-sequencing datasets from cohorts of untreated and matched drug-exposed clinical samples (n=51) revealed that the NCSC gene expression signature (Figure S1A) was also enriched in about 20% of the samples exposed to RAF/MEK-inhibitors when compared to matched treatment naïve counterparts (Figure 1B, S1B and TableS1). Consistently, an increase in cells positive for both NCSC markers, AQP1 and NGFR, could be detected in 2 out of 10 drug-exposed clinical samples analysed. Importantly, these cells were only detected in samples that exhibited a partial response (PR), but absent from samples that did not respond, to the RAF/MEK-inhibitor combinations (Figure 1C). Together, these data confirmed that the NCSC population emerges in about 20% of ON-treatment (O/T) clinical samples and raised the possibility that these cells may function as the cellular origin of nongenetic tumour recurrence.

Testing this possibility requires the ability to perform in-depth longitudinal analyses, as well as intervention and mechanistic studies. We therefore sought to use PDXs as convenient and relevant *in vivo* pre-clinical models. We first probed the diversity of escape mechanisms to a combination of BRAF (i.e. dabrafenib, D) and MEK (i.e. trametinib, T) inhibitors in cohorts of BRAF-mutant PDXs, all established from treatment naïve patients. The treatment inhibited tumour growth to various degrees in almost all PDX models tested (n=9), but eventually all mice progressed ON treatment, confirming that this combination therapy in BRAF-mutant melanoma is limited by the emergence of resistance (Figure 2A and Figure 2B). Strikingly, although the response rates and median survivals varied considerably between models they were, by and large, comparable between mice of the same model. In fact, two distinct behaviours were observed. Whereas drug responses were relatively modest and limited in time for MEL005, MEL007, MEL017, MEL029 and MEL037 (group 1), those of MEL003, MEL006, MEL008 and MEL015 were more marked and tumour regrowth only observed after extended periods of drug-tolerance (group 2). These distinct response curves are in keeping with the various clinical behaviours observed upon long-term follow up of BRAF-mutant melanoma patients^31^. Whereas group 1 reflected what is commonly referred to as intrinsic resistance group 2 mimicked acquired resistance (group 2).

**Figure 2:**
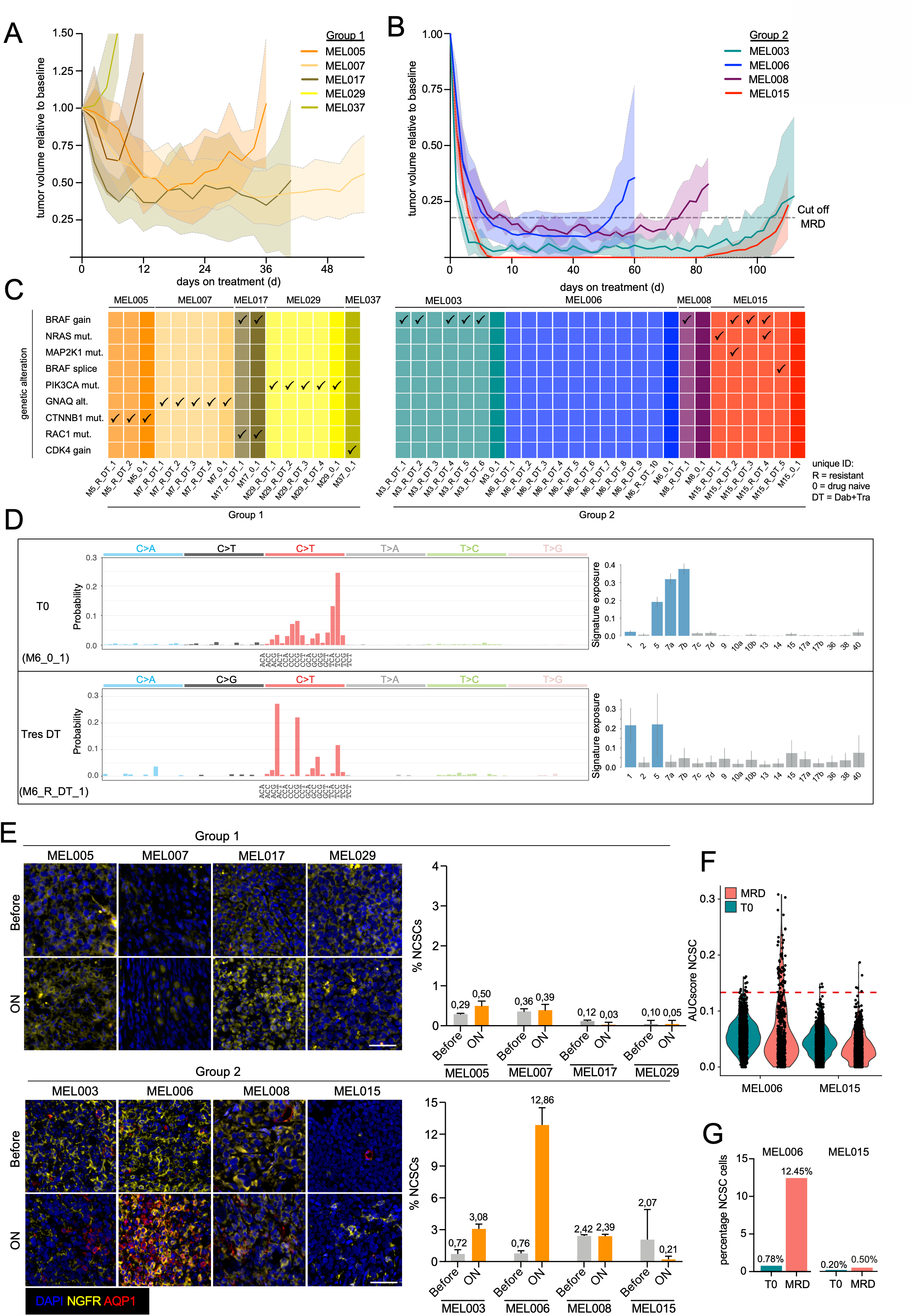
Establishment of in vivo models of nongenetic drug resistance. A) Relative tumour-volume in function of time from Group 1 BRAF-mutant PDXs exposed to BRAF/MEK inhibitors (DT). Group 1; MEL005 (n=6 mice, orange), MEL007 (n=6 mice, light orange), MEL017 (n=6, brown MEL029 (n=6, yellow), MEL037 (n=6, olive green) exposed to DT until resistance. Data is represented by mean (thick line) ±SEM (filled area). B) Relative tumour-volume in function of time from Group 2 BRAF-mutant PDXs exposed to BRAF/MEK inhibitors (DT). MEL003 (n=6 mice) MEL006 (n=18 mice, blue), MEL008 (n=6, burgundy), MEL015 (n=6 mice, red) exposed to DT until resistance. Data is represented by mean (thick line) ±SEM (filled area). C) Interrogation of known genetic driver events of BRAFi&MEKi resistance, including BRAF-mutant splicing variants, in drug naïve lesions (o) and resistant (DT) lesions from BRAF-mutant PDXs using both targeted DNA sequencing and RT-PCR approaches. D) Mutation spectrum of the parental (T0) MEL006 PDX lesion (upper left panel). Mutation types are colour-coded and indicated on top. Trinucleotide contexts are indicated for the C>T mutations and their order is maintained across the other types. COSMIC v3 single base substitution signature exposures for T0 (upper right panel). Sufficiently non-zero signatures are coloured blue. Mutation spectrum of MEL006 PDX lesion (TRes) compared to T0 (lower left panel) and COSMIC v3 single base substitution signature exposures for TRes (lower right panel). E) Immunostainings show the emergence of AQP1/NGFR-double positive NCSCs upon MAPK-inhibition (O/T = ON treatment) in MRD lesions from the MEL006 PDX model. F) Single-cell RNA sequencing of about 20k melanoma cells before treatment (T0) and at MRD showing emergence the NCSC state (based on the AUCell score) in the MEL006, but not MEL015, PDX model. G) Quantification of single cells (from E) harbouring the NCSC-state at T0 versus MRD.

To dissect the mechanisms underlying therapy resistance, we subjected drug naïve (T0) and resistant (TRes) lesions from each of the PDX models from group 1 and 2 to bulk targeted-DNA sequencing analysis. We interrogated 26 loci for the presence of SNAs that are clinically prevalent and/or validated as drivers of resistance to MAPK-therapeutics^23,24,28,29^, and measured *BRAF* copy number. We detected (various combinations of) resistance-conferring genetic events at T0 from all PDXs from group 1 (Figure 2C), indicating that intrinsic resistance in these models is driven by the pre-existence of drug-resistant subclones. In contrast, drug resistance-conferring alterations were absent from all T0 lesions collected from models belonging to group 2. Notably, all resistant tumours collected from the MEL015 cohort^15^ exhibited one or a combination of drug resistance-conferring alterations (Figure 2C). Single-cell targeted DNA sequencing further confirmed the presence of NRAS resistance-conferring mutation in one of the MEL015 TRes samples, but not in the form of rare pre-existing subclone at T0 (Figure S2A). Evidence of *BRAF* amplification was detected in all MEL008 and all but one MEL003 resistant lesions analysed (Figure 2C). These analyses therefore indicated that all these genetic alterations are likely to be acquired *de novo* and were major drivers of the tolerance to resistance switch in these models. Strikingly, the same customized assay did not identify any resistance-conferring event in T0 nor in any of the MEL006 lesions that re-emerged under DT (0/10; Figure 1B), raising the possibility that drug resistance to DT may be systematically driven by nongenetic mechanisms in this PDX model. To rule out the possibility that MEL006 tumours acquire SNAs that are not represented in our targeted screening assay, we performed whole-exome sequencing in matching parental (T0, n=2) and drug-resistant (TRes, n=5) lesions. This analysis did not reveal the presence of any candidate drug-resistance conferring mutations in any of the samples analysed (Figure 1C and data not shown). The vast majority of short variants observed are parental (total 74 indels and 3,860 single and double nucleotide variants) and display the mutational footprint of past UV exposure (COSMIC single base substitution signature SBS7a/b; Figure 1D). In comparison, the TRes lesion accumulated few additional variants (11 indels and 73 single and double nucleotide variants), most of which can be attributed to the activity of the endogenous clock-like mutational processes SBS1 and SBS5 (Figure 1C). Likewise, the allele-specific copy number profiles of T0 and TRes, derived from matching SNP array data, are highly similar, with T0 having one fewer copy of chromosome 7 and an additional copy of chromosome 17 (Figure S2B). While causal drivers of resistance may fail to be identified using the above approaches, selection and expansion of a DT-resistant clone is expected to increase the variant allele frequencies of all passenger mutations present in that clone^32,33^. However, the TRes purity- and copy number-normalised allele frequency distribution (cancer cell fractions), highlights the absence of a single selected, genetically distinct clonal population (Figure S2D). These data demonstrated that resistance to DT in the MEL006, but not MEL015, PDX model systematically resulted from the expansion of a drug-tolerant subpopulation of cells that acquire the ability to grow ON treatment through nongenetic reprogramming. Consistently, resistant lesions from MEL006, but not MEL015, responded to a DT re-challenge following their transplantation and expansion in absence of therapy (drug holiday; Figure S2E). Thus, the MEL006 preclinical model is particularly suited to study the nongenetic mechanisms underlying drug resistance in the relevant *in vivo* context. Together, this analysis confirmed that PDXs are suitable to study the mechanisms underlying drug resistance as the diversity of responses seen in patients is recapitulated in these preclinical models. Unexpectedly, the type of response was very consistent between mice bearing tumours derived from the same patient. Resistance was invariably established through a non-mutational adaptive process in MEL006 and acquisition of *de novo* genetic alterations in MEL015. This observation raises the intriguing possibility that whether resistance occurs through genetic or nongenetic mechanisms may be patient-dependent, deterministic and therefore potentially predictable.

The clinical data reported in Figure 1 pinpointed the NCSC population as the putative cellular origin of nongenetic relapse. Consistently, the NCSC population was either poorly or not detected in lesions from group 1 PDXs, which exhibited intrinsic resistance to MAPK-therapy (Figure 2E and Figure S2F). Immunohistochemistry (IHC) analysis also failed to detect enrichment of NCSCs’ markers in MRD from all three models in which resistance consistently occurs through *de novo* acquisition of genetic alterations (MEL015, MEL003 and MEL008). Note that we only score cells that were positive for both NCSC markers, AQP1 and NGFR. NGFR-positive/AQP1-negative cells could be detected in various lesions both before or ON-treatment, but their presence did not correlate with response. Strikingly, DT-exposure led to a robust increase in the NCSC population in lesions from MEL006 (Figure 2F and Figure S2C). This increase in NCSCs in MEL006, but not MEL015, MRD lesions was further confirmed by single-cell RNA sequencing (scRNA-Seq) experiments (Figure 2F-G).

We therefore reasoned that the cellular composition of MRD may be a key deterministic factor in the decision to engage a genetic versus nongenetic mechanism of resistance, and hypothesized that the presence of the NCSCs, a cell population with increased stem-cell properties (Figure S2G), may favour nongenetic resistance. To test this possibility, we needed to develop an efficient pharmacological approach that fully eradicates these cells from MEL006 MRD. We therefore searched for NCSC-specific molecular vulnerabilities. Gene Set Enrichment Analysis (GSEA) of scRNA-Seq data from individual drug-tolerant cells present in MRD of MEL006 PDX^6^ indicated that Focal Adhesion Kinase (FAK) signalling is the most significantly enriched gene expression signature in the NCSC population (Figure 3A) using the KEGG database as GSEA-reference^34^. This activation appeared specific as it was not detected in the other two drug-tolerant states present in MEL006 MRD (i.e. SMCs and hyperdifferentiated cells; Figure S3A). IHC of MEL006 MRD confirmed selective expression of the activated/phosphorylated form of FAK (pFAK) in geographically localized clusters of NCSCs, which are defined here as double positive for the NCSC-markers NGFR and AQP1 (Figure 3B-C). Cells positive for GFRA2, another NCSC discriminative marker^6^, were then isolated by FACS from an *in vitro* culture of MEL006 cells exposed to DT (Figure 3D). High phosphorylated levels of pFAK were detected by western blotting GFRA2-positive cells. Note that the FAK signalling cascade is activated in many cancers as it confers a cellular proliferative and/or survival advantage by inducing, among others, activation of the AKT pathway in cancer stem cells^35,36^. Accordingly, levels of AKT phosphorylation were elevated in GFRA2-positive cells (Figure 3D).

**Figure 3:**
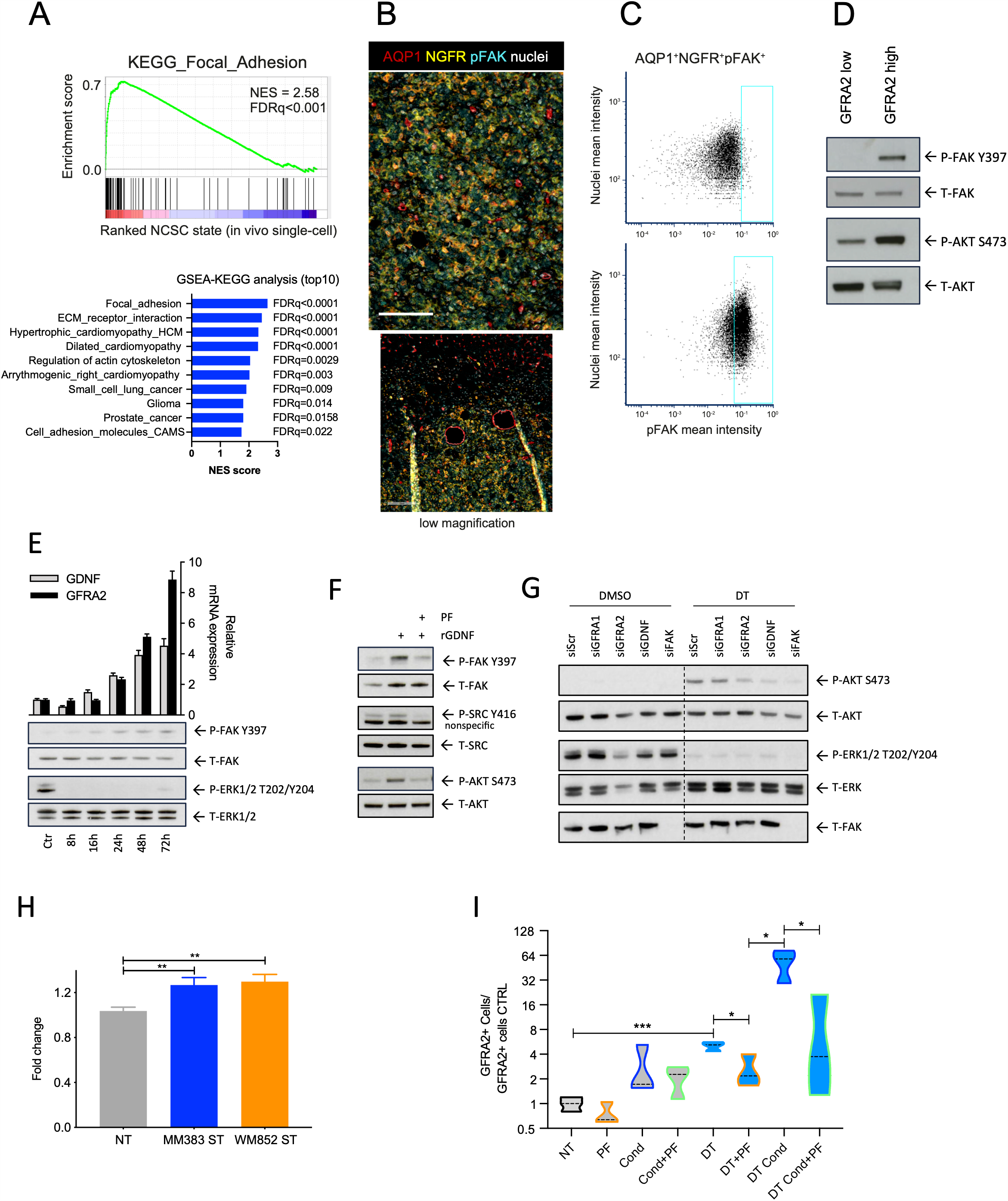
FAK signalling is selectively activated in NCSCs. A) Gene set enrichment analysis (GSEA), using the KEGG as a reference data base, identifies Focal Adhesion expression program as the most enriched in the NCSC state. NES, normalized enrichment score; FDR, false discovery rate. B) Multiplex immunostaining for AQP1 (red), NGFR (yellow), pFAK (cyan), nuclei (white) at minimal residual disease (MRD) following dabrafenib-trametinib (DT) treatment. Representative high (top panel) Scale bar=100µm and low (bottom panel) magnification images are shown Scale bar=500µm. C) Representative image cytometry plots of pFAK intensity in AQP1^-^NGFR^-^ and AQP1^+^NGFR^+^ cells in MEL006 MRD lesions. D) Western blot analysis of total and phosphorylated levels of FAK and AKT in flow-sorted GFRA2^LOW^ and GFRA2^HIGH^ MEL006 melanoma cells. E) Time series monitoring induction of FAK phosphorylation (and ERK inactivation) following DT (100nM/20nM) exposure of an MM383 melanoma culture. Control samples were treated with DMSO. Top graph shows RT-qPCR analysis of *GDNF* and *GFRA2* expression at the indicated time points. Bottom panel shows a representative Western blot analysis of the levels of pan-FAK, phosphorylated FAK at the autophosphorylation site (position Y397), pan-ERK1/2 and phosphorylated-ERK1/2. F) Western blot analysis of overnight serum-starved MM383 cells stimulated with 100ng/ml rGDNF for 1h in the presence (2h pre-treatment before stimulation) or absence of 1uM FAKi PF562271 (PF). Levels of pan- and phosphorylated FAK, AKT and SRC are shown. G) siRNA (80nM) knockdown of *GFRA1, GFRA2, GDNF* and *FAK* in MM383 before and following inhibition of MAPK using DT (100nM/20mM). Western blot analysis shows activation status of AKT and ERK signalling. H) Quantification of GFRA2/NGFR-double positive NCSCs by flow cytometry analysis in non-treated MEL006 cells (NT) and MEL006 cultured in medium supplemented with concentrated supernatant from cultures of MM383 and WM852 cells, containing a high proportion of NCSCs. Error bars represent SD of three biological replicates; *p < 0.05 Mann-Whitney test. I) Quantification of GFRA2+ cells by flow cytometry in short-term cultures from the undifferentiated melanoma cell line MM099 (grey) before (NT) and after 10 days of treatment with 500nM FAKi PF562271 (PF), conditioned medium from MM383 cells (Cond) and conditioned medium plus FAKi (Cond+PF). The experiments were repeated in the presence of BRAF/MEK inhibition, dabrafenib (20nM) and trametinib (4nM) (DT) (blue).

Elevated integrin β1/FAK/Src signalling in melanoma cells was shown to result from paradoxical activation of melanoma-associated fibroblasts by the BRAF-inhibitor and the promotion of matrix production and remodelling^37^. However, FAK activation in NCSCs following MAPK-inhibition did not appear to be strictly dependent on extrinsic factors. DT exposure indeed induced phosphorylation of FAK in BRAF^V600E^-mutant melanoma cultures that exhibit a stable NCSC gene expression profile (i.e. MM383 cell line; Figure 3E and Figure S3B-C). As expected, ERK activation was strongly inhibited by this treatment (Figure 3E). Notably, drug exposure triggered a further increase in the expression of GFRA2 and of another NCSC marker, GDNF (Figure 3E). This observation indicated that MAPK-inhibition triggers FAK signalling in NCSC melanoma cells in a cell-autonomous manner.

GFRA2 is a transmembrane receptor of the GDNF family ligands (GFLs) and an essential transducers of GFLs-mediated activation of signalling pathways that promote survival of several neuronal populations in the central and peripheral nervous system^38,39^. The survival FAK and PI3K-AKT pathways are among the pathways activated by GFRA2-dependent GFLs-mediated signalling^38,39^. As expected the NCSC marker GDNF, one of the GFRA ligands, is expressed in GFRA2^high^ cells (Figure S3B), raising the possibility that FAK activation in NCSCs may be engaged by a GFRA-dependent autocrine loop. Consistently, increasing levels of GDNF were measured in the culture medium of MM383 exposed to DT (Figure S3D). Strikingly, addition of recombinant GDNF alone to the culture medium of MM383 was sufficient to trigger FAK activation and its downstream targets SRC and AKT, even in absence of DT (Figure 3F). Treatment with a pharmacological inhibitor of FAK (PF562271, referred thereafter as PF) confirmed the epistatic relationship with FAK and downstream targets SRC and AKT (Figure 3F). Moreover, silencing of GFRA2, GDNF and FAK by siRNA compromised DT-induced AKT activation in these cells (Figure 3G). In contrast, silencing of another GDNF receptor GFRA1, the expression of which is not induced by DT, did not compromise DT-induced AKT activation. These findings indicated that NCSCs exposed to MAPK-inhibitors are capable of engaging the AKT survival pathway in an autocrine fashion through a GDNF-GFRA2-FAK/SRC signalling cascade.

Interestingly, exposure of an *in vitro* culture of MEL006 PDX to conditioned medium from two distinct NCSC cell lines, the BRAF-mutant MM383 and NRAS-mutant WM852, led to an upregulation of the number of cells positive for the NCSC markers GFRA2 and NGFR (Figure 3H and Figure S3E). We had previously shown that undifferentiated melanoma cells, such as MM099, are capable of switching ON the NCSC phenotype upon concurrent BRAF and/or MEK-inhibition and that therapy-induced emergence of NCSCs can be attenuated by using the RXR-signalling antagonist HX531 (HX)^6^. Consistently, activation of FAK (and SRC) and its downstream target AKT was observed in MM099 exposed to DT and this was attenuated by HX and a pharmacological inhibitor of FAK (PF; Figure S3F). Importantly, an increase in GFRA2-positive cells was also observed following exposure of these cells to MM383 conditioned media (Figure 3I). This effect was exacerbated when the conditioned media was collected following exposure of MM383 cells to DT, but not when cells were exposed to a DT/PF combination. These data indicated that the NCSC transcriptional program can be propagated in a paracrine fashion.

Importantly, a significant correlation was observed between the activities of both FAK and RXR signalling pathways and the NCSC gene expression signatures in large clinical cohorts, highlighting the clinical relevance of the above-findings (Figure S3G).

These data indicated that pharmacological inhibition of FAK signalling may offer a clinically-compatible therapeutic avenue to block emergence of NCSCs in MRD. Small molecule FAK inhibitors have indeed been developed and showed anti-tumour efficacy in several pre-clinical studies, including melanoma^37,40^, and limited adverse effect in patients^41^. Accordingly, several FAK-inhibitors are currently being evaluated in clinical trials across a range of malignancies^41^. Interestingly, exposure to two different FAK inhibitors, PF and defactinib, diminished the drug-dependent emergence of GFRA2-high cells in the MEL006 *in vitro* culture system (Figure 4A and Figure S4A). Moreover, a significant decrease in growth and concomitant increase in apoptotic cell death was observed in GFRA2-high, but not GFRA2-low, cultures upon exposure to DT/PF (Figure 4B-C). Exposure to zVAD, a pan-caspase inhibitor, significantly reversed this effect (Figure 4C). Thus, inhibition of FAK signalling compromises the viability of NCSCs exposed to MAPK-inhibitors *in vitro*.

**Figure 4:**
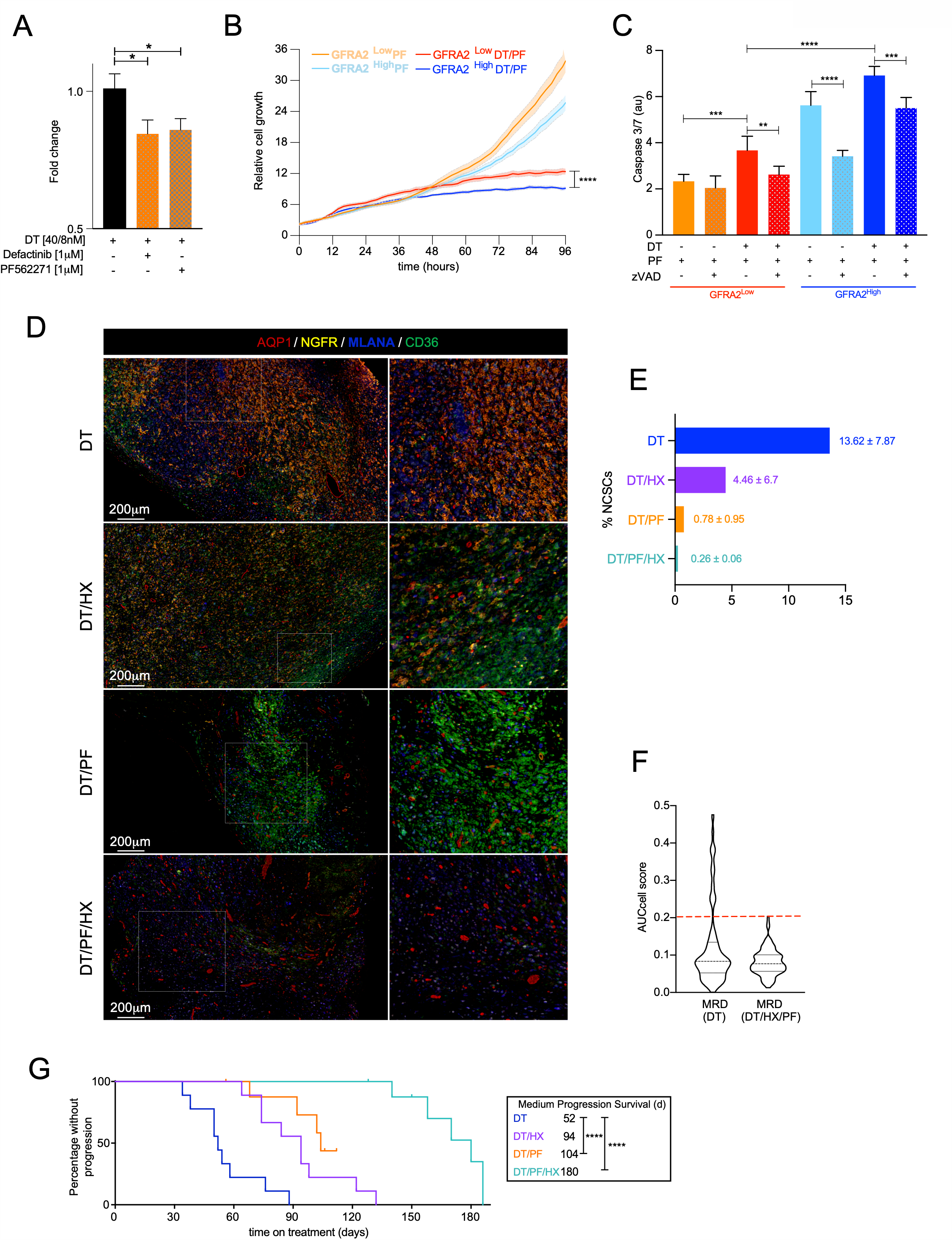
Targeting the NCSCs delays the onset of therapy resistance. A) Quantitative analysis of the number of cells expressing the NCSC markers GFRA2 and NGFR by flow cytometry of MEL006 cells exposed to DT (40nM/8nM) alone or in combination with two FAK inhibitors; Defactenib and PF562271. Error bars represent SD of three biological replicates; *p < 0.05 Mann-Whitney test. B) Relative growth of cultures of GFRA2^high^ (orange) and GFRA2^low^ (Blue) cells sorted by FACS from a MM383 cell culture and exposed to a FAK inhibitor alone (1µM PF562271, PF) (light colour) or in combination with BRAF/MEK inhibitors (40nM Dabrafenib, 8nM Trametinib (DT/PF) (dark colour). The analysis was performed using the IncuCyte live imager. Data (n=6) shows mean (thick line) ±SEM (filled area). C) Caspase 3/7 activity in cultures of GFRA2^high^ (orange) and GFRA2^low^ (Blue) cells sorted by FACS from a MM383 cell culture and exposed to a FAK inhibitor alone (1µM PF562271, PF) (light colour) or in combination with BRAF/MEK inhibitors (40nM Dabrafenib, 8nM Trametinib (DT/PF) (dark colour) and a pan-*caspase* inhibitor, zVAD (50mM) (polka dot pattern). Data show mean of biological replicates (n=6) ±SD. D) Multiplex immunostaining for AQP1 (red), NGFR (yellow), MLANA (blue) and CD36 (green) in minimal residual disease (MRD) following combination treatment with dabrafenib-trametinib (DT), an RXR antagonist (HX), and focal adhesion kinase inhibitor (PF). E) Quantification of AQP1/NGFR-double positive NCSC cells from multiplex immunostaining (D). The frequency of NGFR/AQP1-double positive cells are indicated as percentage per area ±SEM (n=3 biological replicates; 3 areas per sample). Scale bar=200µm. F) Comparison of the cellular composition of MRD from MEL006 lesions treated with DT versus DT/HX/PF using single-cell RNA sequencing. The NCSC activity per cell was inferred using AUCell (Aibar et al. 2017). NCSC cells were detected from DT, but not DT/PF/HX, MRD (AUCell score <0.2). G) Kaplan-Meier curve for MEL006 mice treated with dabrafenib-trametinib alone (DT, n = 9)^6^, DT plus HX531 (DT/HX, n = 9)^6^, DT plus FAK inhibitor, PF562271 (DT/PF, n = 9) and the quadruple combination (DT/PF/HX). Median time to progression was 52 days for DT, 94 days for DT/HX, 104 days for DT/PF and 180 days for DT/PF/HX. Log rank (Mantel-Cox) for DT versus DT/PF: p < 0.0001 (****) and DT versus DT/PF/HX: p < 0.0001 (****).

Given the important contribution of NCSCs in the development of therapy resistance, combining PF with MAPK therapeutics may significantly limit the risk of relapse. To test the therapeutic potential of this regimen, we first assessed the impact of PF on the cellular composition of MEL006 MRD. We performed IHC analyses on MRD materials from mice treated with DT and DT/PF. While, as expected, an increase in NCSC markers was observed in MRD isolated from mice treated with DT, this was dramatically attenuated upon exposure to the triple combination (DT/PF; Figure 4D-E). Likewise, and consistent with our previous study^6^, exposure to HX also decreased the therapy-induced emergence of NCSCs at MRD. These decreases resulted in an increase in CD36- and MLANA-positive cells. Strikingly, the number of NCSCs present in MRD from mice treated with the quadruple combination DT/PF/HX dropped even further, so much so that none could be detected in most lesions analysed. Histological analysis (H&E staining) showed a progressive increase in highly pigmented (hyperdifferentiated) melanoma cells from MRD lesions exposed to DT, DT/PF and DT/PF/HX and a concomitant decrease in tissue integrity due to necrosis and edema (Figure S4B). A scRNA-seq experiment confirmed the absence of cells harbouring the NCSC gene expression signature in MRD lesions collected from mice treated with the DT/PF/HX combination (Figure 4F and Figure S4C).

Importantly, PF administration increased the response rates to the DT or even the DT/HX combination (Figure S4D). Moreover, combining PF with DT or DT/HX produced a significantly longer median progression-free-survival (PFS) period compared to the DT treatment and delayed the development of resistance (Figure 4G; Median PFS for DT=52days, DT/HX= 94days, DT/PF= 104 and DT/PF/HX=180 days). Importantly, such treatments did not cause any relevant adverse reaction or weight loss (Figure S4E). It is noteworthy that, although the DT/PF combination was slightly superior than DT in delaying tumour recurrence in the MEL015 PDX model, the effect was far less pronounced than in the MEL006 cohorts (Figure S4E). These data identified a clinically-compatible methodology that efficiently prevents the accumulation of the drug-tolerant NCSC population in MRD and further confirmed that this subpopulation is a major driver of therapy resistance.

The ability to efficiently abrogate the emergence of NCSCs in an *in vivo* clinically-relevant setting gave us the opportunity to test whether transition from drug tolerance to resistance depends on a particular cellular MRD composition. We subjected all resistant lesions to bulk targeted-DNA sequencing analysis. While as described above, we failed to identify resistance-conferring events in lesions that re-emerged under DT (0/10), resistance-conferring genetic alterations were identified in most (8/9) MEL006 lesions that acquired resistance to the DT/PF/HX combination (Figure 5A). One of these lesions (M6_R_DTHXPF_04) carried a NRAS^Q61K^ activating mutation, previously described to confer resistance to BRAF/MEK-inhibition^42^ (Figure S5A). *BRAF* amplification, a particularly common resistance-conferring event^30^, was detected in most (7/9) DT/PF/HX-resistant lesions, but not in any of the T0 or DT-resistant lesions. A quantitative-PCR DNA analysis confirmed the increase in *BRAF*, but not *CRAF*, copy number in those samples (M6_R_DTHXPF_01, 02, 06, 08 and M6_R_DTHXPF_09), but not in DT resistant lesions analysed (Figure 5B and Figure S5C). The increase in *BRAF* copy number was further confirmed at single-cell resolution by DNA-FISH (Figure 5C and Figure S5B). This latter analysis showed that the vast majority of cells carried the *BRAF* gene amplification, consistent with a clonal cell population.

**Figure 5:**
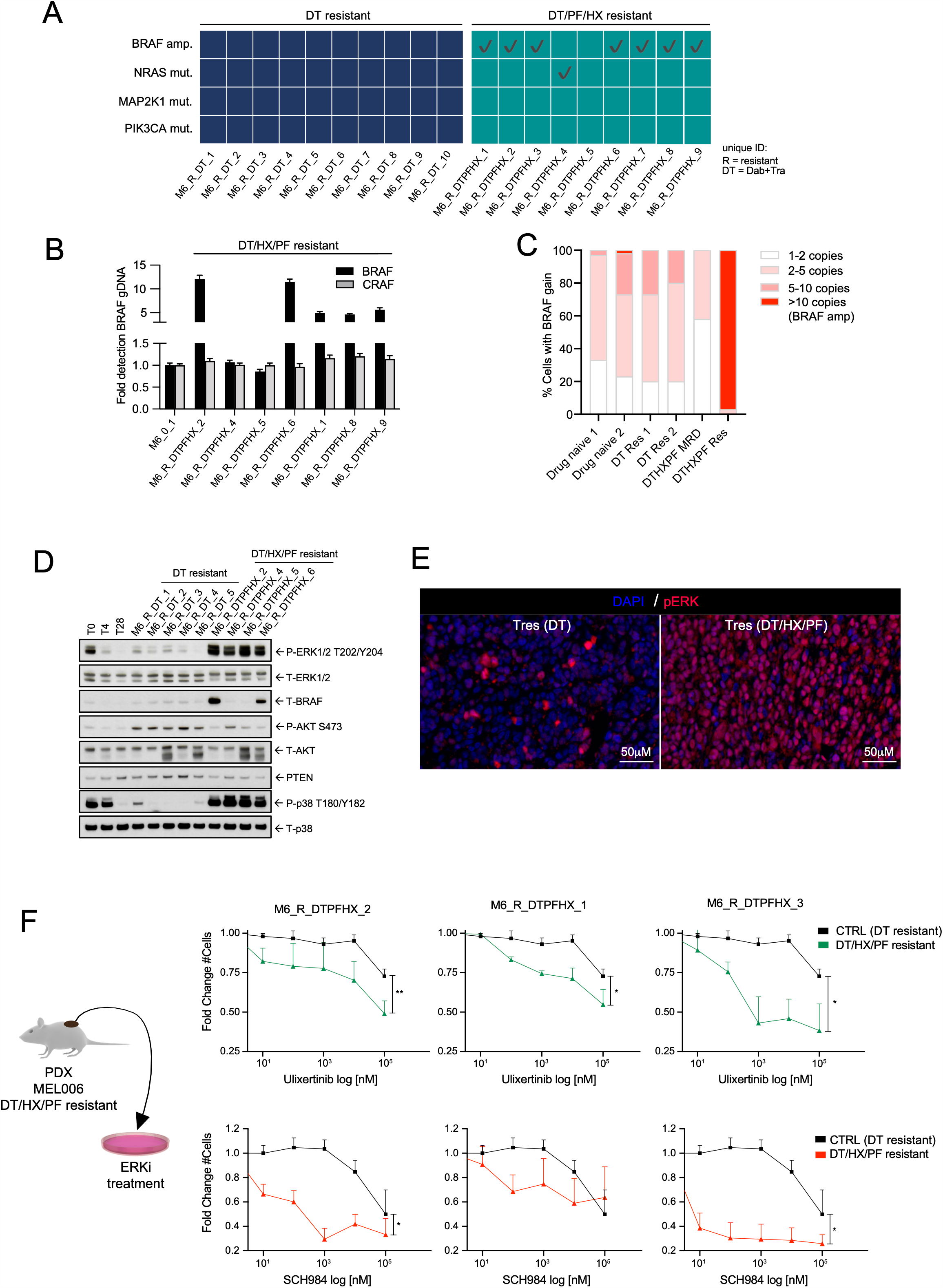
Targeting the NCSCs prevents nongenetic drug resistance evolution. A) Interrogation of known genetic driver events of BRAFi&MEKi resistance, including BRAF-mutant splicing variants, in MEL006 lesions that acquired resistance to DT or DT/PF/HX lesions using both targeted DNA sequencing and RT-PCR approaches. B) Quantitative PCR analysis for *BRAF* and *CRAF* copy number was performed from genomic DNA. (RQ=relative number of copies compared to T0). C) Quantification of percentage of cells exhibiting *BRAF* copy gain as assessed by DNA FISH analysis using Dual Colour Break Apart Probe (ZytoVision) on PDX tissue sections from treatment naïve (T0) and lesions that acquired resistance to DT or DT/PF/HX. D) Western blot analysis of levels of ERK and AKT phosphorylation and BRAF protein expression in lysates from MEL006 PDX tumours before treatment (T0) and ON treatment with DT for 4 days (T4), at MRD (T28) and after the development of resistance to DT or DT/HX/PF. E) Representative immunofluorescence staining with anti-p-ERK antibodies (red) in resistant lesions that escaped the DT (Tres DT01 is shown) and DT/HX/PF (qTRes01 is shown) treatments. Slides were counterstained with DAPI (blue). F) Cell viability assay of MEL06 primary cells (CTRL) and primary cells derived from MEL06 DTPFHX resistant lesions harbouring *BRAF* gene amplification (M6_R_DTPFHX_1, M6_R_DTPFHX_2 and M6_R_DTPFHX_3) treated with ERK inhibitors (Ulixertinib, green and SCH984, red). Mean (n=4); bars, SD; * p < 0.05 **, p < 0.01.

We failed to detect a genetic cause of resistance in only one (M6_R_DTHXPF_05) of the 9 lesions analysed. It remains unclear whether resistance in this case was driven by a genetic alteration that we did not screen for or through a nongenetic mechanism. A western blot analysis of this sample, as well as the NRAS-mutant (M6_R_DTHXPF_04) and two BRAF-amplified samples (M6_R_DTHXPF_02 and 06), showed a similarly dramatic elevation in p-ERK (and p-P38) levels (Figure 5D-E). In comparison, levels of p-ERK (and p-P38) were much lower in DT-resistant lesions (5/5), all of which exhibited increased p-AKT levels instead. Although more extensive genetic analysis is required to draw firm conclusions, this observation indicates that resistance in M6_R_DTHXPF_05 might also have a genetic origin. More importantly, the data also raise the possibility that whereas nongenetic drug resistance evolution may favour activation of AKT signalling, genetic mechanisms seemingly prioritize ERK reactivation.

We therefore reasoned that lesions that escaped the NCSC-directed therapy through acquisition of genetic alterations maybe sensitive to ERK-inhibition. Accordingly, exposure of cells from three different DT/HX/PF MEL006 resistant lesions to increasing concentrations of two different ERK inhibitors showed that the resistant cells were far more sensitive than their matching control cells (Ctr; Figure 5F). This observation therefore provides a clinically-viable approach for the treatment of patients that would progress on NCSC-targeted therapies.

Notably, immunostaining demonstrated that the increase in p-ERK levels was homogeneously distributed among virtually all individual cells in DT/HX/PF (but not DT) resistant tumours, again consistent with resistance being driven by a clonal genetic event in the DT/HX/PF resistant lesions (Figure 5D-E). Because exposure to PF and HX eradicated emergence of DT-induced NCSCs (Figure 4C-D), these data indicated that the presence of the NCSC drug-tolerant subpopulation at MRD is required for development of nongenetic resistance. Together, these data further establish a strict correlation between the presence of NCSCs at MRD and the development of therapy resistance through nongenetic mechanisms.

## Discussion

We have previously shown that transient de-differentiation of melanoma cells into a neural crest stem cell (NCSC) state contributes to the development of resistance to targeted therapy^6^. One of the key findings we report herein is that the NCSCs rely on FAK-signalling for growth and survival. This parallels a recent observation that a neural crest stem cell population, which contributes to jaw bone regeneration, gains activity within the FAK signalling cascade and that inhibiting FAK abolishes new bone formation^43^. It is also related to some degree to another study, which reported increased FAK signalling in melanoma cells exposed to a BRAF-inhibitor^37^. This study, however, reported that activation of FAK in melanoma cells is driven by a “paradoxical” activation of melanoma-associated fibroblasts, induction of matrix production and remodelling leading to elevated integrin β1/FAK/Src signalling^37^. In contrast, we show herein that therapy-induced FAK signalling in the NCSC population is caused by the activation of an autocrine loop in which GDNF, produced by the NCSCs themselves, engages GFRA2-dependent activation of FAK signalling and one of its downstream pro-survival targets, AKT. Interestingly, we also show that NCSCs can promote activation of the NCSC transcriptional program in a paracrine fashion, an observation that may, at least partly, explain why NCSCs tend to occur in geographically localized clusters in MRD lesions^6^. The molecular mechanisms underlying this paracrine effect remains to be elucidated. GDNF, which is produced by the NCSCs, may act as an inducer of phenotypic conversion into a NCSC state through its ability to induce signalling independently of GFRalpha, for example via the RET or MET receptors^44^. These receptors are indeed expressed in melanoma cells, including those that do not harbour a NCSC state, and their activation by GDNF may indeed initiate such a phenotypic switch. In keeping, GDNF was shown to contribute to paracrine regulation in a multitude of biological processes, such as for example spermatogonial self-renewal and differentiation^45^. It will therefore be interesting to test whether GDNF (or other factors) secreted by the NCSCs is key trigger of this paracrine.

The observation that NCSCs rely on FAK signalling has several important clinical implications. We show that exposure to FAK-inhibitors strongly decreases emergence of the NCSCs in MRD lesions, and drastically delays the onset of resistance to RAF/MEK inhibitors in preclinical PDX models. Moreover, we show that although emergence of the NCSC population in MRD is strongly compromised upon FAK-inhibition, it can only be fully abolished by combining PF with HX, an RXR antagonist. Accordingly, this regimen produced a significantly longer median Progression Free Survival (PFS) period than the DT/PF combination and even further delayed the development of resistance. Note that this treatment did not cause any relevant adverse reaction or weight loss in PDXs. Our study therefore provides a strong rationale for the testing of the DT/PF and DT/PF/HX combinations in BRAF^V600E^-mutant melanoma patients. Moreover, FAK physically and functionally interacts with Src to promote a variety of cellular responses and accordingly we find a strong correlation between FAK/Src expression/phosphorylation in NCSCs upon DT exposure. Anticancer agents that target Src are also under development in a broad variety of solid tumours^46^. It may therefore also be interesting to further test combinations of both FAK and Src inhibitors, such as Dasatinib, as a more effective strategy to suppress the emergence of the NCSC population at MRD.

Importantly, de-differentiation into a NCSC-like state was also reported as an escape mechanism to T-cell transfer therapy^47-49^. This can be explained by the drastic decrease of expression of melanoma antigens, including MART1/gp100, as cells de-differentiate^50,51^. Because NCSCs exhibit stem cell features, we had postulated that this subpopulation may also activate mechanisms allowing them to escape the immune defence and, thereby, provide a pool of cells that are refractory to immune checkpoint blockade (ICB)^13^. The presence of these cells in tumours before exposure to ICB or their accumulation during treatment may therefore contribute, respectively, to both intrinsic and adaptive resistance to immunotherapy. Together these findings highlighted the urgent need for therapeutic strategies directed at this melanoma subpopulation and emphasized how wide is the fraction of melanoma patients that may benefit from (the above-described) NCSC-directed therapy. Importantly, although pharmacological eradication of the NCSC population in PDXs was sufficient to avoid the development of nongenetic resistance it did not prevent relapse. Resistance eventually developed through acquisition of *de novo* resistance-conferring genetic alterations. The prediction from these experiments is that, although useful to delay PFS, NCSC-directed therapies are unlikely to be curative. However, these experiments were conducted on an immunocompromised background (i.e. nude mice) and MRD lesions from mice exposed to the DT/PF/HX combination were almost exclusively composed of hyperdifferentiated melanoma cells. It remains possible that these cells may eventually be cleared by the immune system. Moreover, these cells are likely to be highly responsive to both T-cell therapy or ICB. These data therefore also warrant the implementation into the clinic of a sequential treatment regimen in which tumour debulking is induced by the DT/PF/HX combination followed by an immunotherapy approach aiming at eradicating the residual pool of hyperdifferentiated cells.

It has been proposed that discrete subpopulations of cancer cells may harbour increased “epigenetic” plasticity that permits random activation of alternate gene regulatory networks and, thus allows acquisition of specific phenotypic properties through nongenetic reprogramming^26^. Some of these properties may be maintained through cell division, and eventually lead to the selection of drug resistant “epiclones”. Consistently, the findings described herein identify the NCSCs as a melanoma subpopulation that is highly permissive to nongenetic reprogramming and a key driver of nongenetic resistance. Interestingly, we also provide evidence that nongenetic resistance can develop in absence of selection of a single, genetically-distinct, clonal population, a conclusion that is further supported by the recent re-analysis of our single-cell data with LACE (Longitudinal Analysis of Cancer Evolution), an algorithm that processes single-cell somatic mutation profiles from scRNAseq data^52^. Together, these analyses raise the possibility that nongenetic resistance may not be clonal but develop, instead, through collective reprogramming.

The ability in PDX models to repeatedly treat the same tumour over and over again led to the key and rather unexpected finding that a given tumour recurrently selects either a genetic or nongenetic drug resistance trajectory. We provided evidence that nongenetic resistance only develops following emergence of the NCSCs in MRD. These data indicate that the cellular composition of MRD dictates whether resistance develops through a genetic or nongenetic mechanism and therefore that this process is deterministic and predictable. The clinical implications of this finding are far-reaching. For instance, whether a given patient will (or not) benefit from combination treatments that make use of epigenetic drugs (for which several clinical trials are ongoing) will depend on the MRD composition. Given the inter-patient variability of the cellular composition of MRD, our data highlight the need to develop personalized MRD-targeting therapies. Note that the clinical implementation of such therapies will require the ability to access serial tumour samples and their deep analysis using, for instance, emerging single-cell spatial multi-omics methods. Because access to ON-treatment biopsies is often problematic in the context of solid cancers, such as melanoma, a possible future alternative will be the development of non-invasive and ultrasensitive methods able to capture cellular composition of MRD from liquid biopsies. We argue that priority should be given to these technological developments as they will likely lead to promising therapeutic avenues that negate the increasingly-recognized contribution of non-genetic mechanisms to therapy resistance.

## MATERIAL and METHODS

### Drugs

BRAFi Dabrafenib, MEKi Trametinib and FAKi PF562271 were purchased from MedChemExpress, RXR antagonist HX531 from Tocris, siRNA SMARTpools from Dharmacon (siRNA sequences are available upon request), recombinant GDNF from Peprotech.

### Tissue Culture

Cells from dissociated MEL006 tumours and MM099 short culture melanoma cells were grown in 5% CO2 at 37°C in F10 supplemented with 10% FBS, 0.25% GlutaMAX and Penicilline/Streptomycin antibiotics. MM383 and WM852 cells were grown in 5% CO2 at 37°C in RPMI GlutaMAX supplemented with 10% FBS and Penicilline/Streptomycin antibiotics.

### Apoptosis assay

IncuCyte® Caspase-3/7 Red Apoptosis Assay Reagent (#4704) were used to measure apoptosis on an IncuCyte ZOOM system (Essen BioScience). The red fluorescence of 500 seeded cells on 96-well plate (TPP) were automated measured on pictures taken at 2 h intervals for the duration of the experiments.

### Western blotting

Harvested cell culture pellets were resuspended in protein lysis buffer (25mM HEPES pH 7,5; 0,3M NaCl; 1,5mM MgCl2; 2mM EDTA; 2mM EGTA; 1mM DTT; 1% Triton X-100; 10% Glycerol; phosphatase/protease inhibitor cocktail), incubated on ice (15min) and centrifuged (15min) at 4°C/13000 rpm. Tumour samples were additionally homogenized with a PreCellys in protein lysis buffer, prior to incubation on ice. Equal amounts of protein (Bradford quantification) were run on 4-12% Bis-Tris Plus Bolt gels (ThermoFisherScientific) and transferred to a nitrocellulose membrane with an iBlot dryblot system (ThermoFisherScientific). Membrane blocking (5% BSA/TBS-0,2%Tween) is followed by incubation with the appropriate primary antibodies and HRP-conjugated secondary antibody (Cell Signaling). Signals were detected by enhanced chemiluminescence (ThermoFisherScientific) on Amersham hyperfilm. Antibodies were from Cell Signaling Technology (P-FAK Y397, #8556; FAK, #13009; P-SRC Y416, #6943; SRC, #2123; P-AKT S473, #4060; AKT, #4691; P-ERK T202/Y204, #9106 (cell lines); P-ERK T202/Y204 #4370 (tumour lysates); ERK, #9102; BRAF, #9433; P-p38 T180/Y182, #4511; p38, #8690; PTEN, #9559)

### RT-qPCR

Cells were harvested and mRNA extracted using the RNA NucleoSpin extraction kit (Macherey&Nagel). RNA was quantified using a NanoDrop 1000 (Thermo Scientific) and 500–2,000 ng was reverse transcribed with a High-Capacity cDNA Reverse Transcription Kit (ThermoFisherScientific). qPCRs were run using SensiFAST SYBR No-ROX Kit (Bioline) and a Roche LightCycler 384. Data processing with qbase+ 3.1 software (Biogazelle) relies on normalization with a minimum of 2 reference genes. RT-qPCR primer sequences are available upon request.

### Patient-derived xenografts

In collaboration with TRACE, patient-derived xenografts (PDX) models were established using tissue from patients undergoing surgery as part of standard-of-care melanoma treatment at the University Hospitals KU Leuven. Written informed consent was obtained from all patients and all procedures involving human samples were approved by the UZ Leuven Medical Ethical Committee (S54185/S57760/S59199) and carried out in accordance with the principles of the Declaration of Helsinki. PDX models MEL006, MEL015 and MEL029. All procedures involving animals were performed in accordance with the guidelines of the IACUC and KU Leuven and were carried out within the context of approved project applications P038/2015, P098/2015 and P035/2016. Fresh tumour tissue was collected in transport medium (RPMI1640 medium supplemented with penicillin/streptomycin and amphotericin B). Tumour fragments were subsequently rinsed in phosphate-buffered saline supplemented with penicillin/streptomycin and amphotericin B and cut into small pieces of approximately 3 × 3 × 3 mm3. Tumour pieces were implanted subcutaneously in the interscapular fat pad of female SCID-beige mice (Taconic). Sedation and analgesia were performed using ketamine, medetomidine and buprenorphine. After reaching generation 4 (F4), one mouse with a tumour of 1000 mm3 was sacrificed. This tumour was minced followed by dissociation using collagenase I & IV and trypsin. Cells were resuspended in serum-free DMEM/F12 medium and 250 000 cells were injected in the interscapular fat pad of 8 – 16 week old female NMRI nude mice (Taconic). For single cell RNA sequencing purposes, cells were transduced with a lentivirus carrying dsRed. Cells were washed four times before injecting into the interscapular fat pad. For immunohistochemistry, non-dsRed-transduced lesions were used. For FACS, tumours were enzymatically dissociated using the same protocol.

### Pharmacologic treatment of mice

Mice harboring tumours of a comparable size, ranging from 900 to 1000 mm3, were exposed to the BRAF-MEK combination via daily oral gavage. BRAF inhibitor dabrafenib and MEK inhibitor trametinib were dissolved in DMSO at a concentration of 30 and 0.3 mg/mL respectively, aliquoted and stored at −80°C. Each day a fresh aliquot was thawed and diluted 1:10 with phosphate-buffered saline. Mice were treated with a capped dose of 600 – 6 μg dabrafenib – trametinib respectively in 200 μL total volume. For the dabrafenib-trametinib-HX531 and dabrafenib-tramatenib-PF562271 preparation, HX531 at 10 mg/dL and PF562271 at 45mg/mL respectively in combination with dabrafenib and trametinib. Tumour volume was monitored with a caliper and the volume was calculated using the following formula: V = (π/6) x length x width x height.

### Fluorescence *in situ* hybridization analysis

Interphase fluorescence *in situ* hybridization (FISH) was performed on 5-μm paraffin sections of formalin fixed (FFPE) xenograft tumour specimens applying the BRAF SPEC BRAF Dual Color Break Apart Probe (ZytoVision GmbH, Bremerhoven, Germany). Briefly, FFPE sections were deparaffinized in three changes of xylenes, dehydrated in ethanol, pretreated in sodium thiocyanate buffer (Abbott Molecular, Abbott Park, IL) for 30 minutes at 80°C, followed by pepsin digestion for 25 minutes at 37°C. Hybridization was performed at 37°C overnight. Slides were then washed and mounted with DAPI in an antifade solution. The number of fused BRAF signals was analyzed in 100 hundred successive, non-overlapping tumour cell nuclei using a Zeiss fluorescence microscope (Zeiss Axioplan, Oberkochen, Germany), controlled by Isis 5 software (Metasystems, Newton, MA).

### Immunofluorescence on PDX biopsies

Fluorescent staining was performed using OPAL staining reagents, which use individual tyramide signal amplification (TSA)-conjugated fluorophores to detect various targets within an immunofluorecence assay. In brief, samples were fixed at 4% Paraformaldehyde and embedded in paraffin. Serially cut sections of 5 μm were stained with haematoxylin and eosin for routine light microscopy, and used for immunohistochemistry. Depending on the antibody, antigen retrieval was performed in Citrate buffer at pH 6. Deparaffinized sections were then incubated overnight with primary antibodies against AQP1 (cat No. #AB2219, Millipore), NGFR (cat No. 8238, Cell Signalling Technology) and phospho-ERK1/2 Thr202/Tyr204 (Cat No. #4370, Cell Signalling Technology). Subsequently, the slides were washed in phosphate-buffered saline, pH 7.2, and incubated for 10 min at room temperature with Opal Polymer HRP Mouse Plus Rabbit secondaries (PerkinElmer). After another wash in PBS, the slides were then incubated at RT for 10 min with one of the following Alexa Fluorescent tyramides (PerkinElmer) included in the Opal 4 colour kit (NEL810001KT) to detect antibody staining, prepared according to the manufacturer’s instructions: Opal 520, Opal 570 and Opal 690 (dilution 1:50).

Stripping of primary and secondary antibodies was performed by placing the slides in a plastic container filled with antigen retrieval (AR) buffer in Citrate pH 6; microwave technology was used to bring the liquid at 100 °C (2 min), and the sections were then microwaved for an additional 15 min at 75 °C. Slides were allowed to cool in the AR buffer for 15 min at room temperature and were then rinsed with deionized water and 1 × Tris-buffered saline with Tween 20. After three additional washes in deionized water, the slides were counterstained with DAPI for 5 min and mounted with ProLong™ Gold Antifade Mountant (Thermofisher Scientific). Slides were scanned for image acquisition using Zeiss AxioScan Z.1 and ZEN2 software.

### Multiplexed, Sequential Immunohistochemistry and Analysis

Sequential chromogenic immunohistochemistry (IHC) was performed as previously described ^53^. In brief, 5 µm FFPE tissue sections of PDX samples were de-paraffinized, bleached (10% hydrogen peroxide at 65°C, 10 min), and subsequently stained with hematoxylin (GHS116, Sigma-Aldrich). Iterative cycles of standard IHC (heated antigen retrieval, Citra Plus, pH 6.2, BioGenex) were performed followed by detection with ImmPressTM IgG-polymerized peroxidase reagents (Vector Laboratories) and visualization with AEC (Vector Laboratories). Slides were scanned and subsequently, AEC was removed using ethanol and antibody stripped in heated citrate buffer to allow the next staining cycle. Complete antibody removal was confirmed at each step. Images were acquired using Aperio ImageScope AT (Leica Biosystems). Serial digitized images were processed using a computational image analysis workflow described previously^53^ and images aligned and pseudo-coloured for visualization (Aperio ImageScope, Leica). Single-cell segmentation and quantification of staining intensity was performed using a CellProfiler v.2.1.1 pipeline. FCS Express 5 Image Cytometry v.5.01.0029 (De Novo Software) was used to gate intensity thresholds for subsequent analysis. Cell numbers extracted from FCS Express was used to generate image cytometry plots and plot relative proportions.

### FACS

Cells were incubated with GFRA2 (R&D systems, AF429) and NGFR (Cell Signalling Technology, #8238) antibody for 45minutes at room temperature, followed by a secondary antibody conjugated with Alexa Fluor® 594 or Alexa Fluor® 647 for 30 min at room temperature. Cells were resuspended in FACS sorting buffer (culture medium supplied with 5% serum and 2 mM EDTA). FACS analyses were performed with BD FACSChorus™ and FlowJo® software.

### Single cell sorting and SMARTseq2 based scRNA sequencing

Single cell suspensions of MRD (DT/HX/PF) lesions were obtained as previously described^6^. DAPI negative, CD44 positive cells were sorted (BD Influx) in 96 well plates (VWR, DNase, RNase free) containing 2 μL of lysis buffer (0.2% Triton X-100, 4U of RNase inhibitor, Takara) per well. Plates were properly sealed and spun down at 2000 g for 1 min before storing at −80°C. SMART-seq2 based scRNAseq was performed as previously described^6^. NCSC signature activity^6^ per single cell was inferred using AUCell^54^.

### Targeted scDNA sequencing

Nuclei were extracted from a fresh frozen MEL015 treatment-naïve tumour sample according to the manufacturer’s protocol (Tapestri, missionbio). Single-cell sequencing was performed using Mission Bio’s Tapestri THP platform, which assesses hotspot mutations in about 50 genes, according to the manufacturer’s protocol. Nuclei were emulsified with lysis reagent and incubated at 50° C prior to thermally inactivating the protease. The emulsion containing the lysates from protease-treated single-cells was then microfluidically combined with targeted gene-specific primers, PCR reagents, and hydrogel beads carrying cell identifying molecular barcodes using the Tapestri instrument and cartridge. Following generation of this second, PCR-ready emulsion, molecular barcodes were photocleavably released from the hydrogels with UV exposure and the emulsion was thermocycled to incorporate the barcode identifiers into amplified DNA from the targeted genomic loci. The emulsions were then broken using perfluoro-1-octanol and the aqueous fraction was diluted in water and collected for DNA purification with SPRI beads (Beckman Coulter). Sample indexes and Illumina adaptor sequences were then added via a 10 cycle PCR reaction and the amplified material was then SPRI purified a second time. Following the second PCR and SPRI purification, full-length amplicons were ready for quantification and sequencing. Libraries were analysed on a DNA 1000 assay chip with a Bioanalyzer (Agilent Technologies), and sequenced on the Illumina Hiseq2500 platform (PE150bp, rapid mode). Sequencing data were processed using Mission Bio’s Tapestri Pipeline (trim adapters using cutadapt, sequence alignment to human reference genome hg19, barcode demultiplexing, cell-based genotypecallingusingGATK/Haplotypecaller). Data were analysed using MissionBio’s Tapestri Insights software package and visualized using R software.

### Droplet based scRNA sequencing

We profiled 4 PDX melanoma lesions (MEL006: 1xT0, 1xMRD and MEL015: 1xT0, 1xTres) using 3’ scRNA sequencing (10x genomics) with a target cell recovery ranging from 5-10k cells per sample. Libraries for single cell RNA sequencing were constructed using the 10X Genomics Chromium platform according to manufacturer’s instructions. Library construction was primarily done with the Chromium Single Cell 3’ GEM, Library & Gel Bead Kit v3. In brief, cells were partitioned into Gel Bead-In-Emulsions (GEMs) at limiting dilution, where lysis and reverse transcription occurred yielding uniquely barcoded full-length cDNA from poly-adenylated mRNA. GEMs were subsequently broken and the pooled fraction was amplified, followed by fragmentation, end repair and adaptor ligation of size selected fractions. Transcriptome libraries were sequenced with paired end reads on an Illumina NovaSeq6000. After quality control, the raw sequencing reads were aligned to the human reference genome v. GRCh38 and mouse reference genome (mm10), followed by application of CellRanger (10x Genomics, v2.0) in order to obtain feature-barcode matrices. Cells with higher mapping results to the mouse genome were removed as well as potential doublets using the DoubletFinder v. 2.0.2 pipeline. Raw count matrices were analysed using R package Seurat v. 3.1.3. The matrices were filtered by removing cell barcodes with <1000 expressed genes, >9000 expressed genes and >30% (for T0 samples) and >50% (for MRD samples) of reads mapping to mitochondrial genes.

### Targeted bulk DNAseq

DNA extraction from fresh frozen minced PDX tumour tissue was performed using the DNeasy Blood&Tissue kit (Qiagen) according to the manufacturer’s instructions. DNA was quantified using the Qubit 2.0 fluorometer (Life Technologies, Carlsbad, CA, USA). Analysis for hotspot mutations was performed with the TruSight Tumour26 or Tumour97 kit (Illumina, San Diego, CA, USA), which enables the detection of mutations in 26 genes (AKT1, ALK, APC, BRAF, CDH1, CTNNB1, EGFR, ERBB2, FBXW7, FGFR2, FOXL2, GNAQ, GNAS, KIT, KRAS, MAP2K1, MET, MSH6, NRAS, PDGFRA, PIK3CA, PTEN, SMAD4, SRC, STK11, TP53). The TruSight technology is based on extension and ligation-based amplicon library preparation specific for each of the two strands of DNA. The two independent libraries were combined and sequenced on a MiSeq/NextSeq instrument (Illumina) by paired-end sequencing (2×121 bp) with a minimum read depth of at least 1000× coverage, according to the manufacturer’s instructions. The paired-end reads were mapped against the Genome Reference Consortium Human Build 19 (GRCh37). Data analysis was performed using an in-house developed bioinformatics pipeline that begins with the FASTQ files and incorporates BWA for alignment, GATK for variant calling, and Annovar for variant annotation^55-57^.

### Copy number analysis

Genomic DNA was extracted for MEL006 and MEL015 T0 and Tres using the DNeasy Blood&Tissue kit (Qiagen) according to the manufacturer’s instructions. DNA samples were quantified using UV absorbance, and SNPs at ± 300,000 sites were determined using the whole genome scanning 12-sample Illumina HumanCytoSNP-12v2.1 BeadChip according to the manufacturer’s instructions. Images were captured on Cytoscan (Illumina), and data were primarily analysed using Illumina’s GenomeStudio software.

Allele-specific copy number and sample purities were inferred using ASCAT (v2.5.2) after germline genotype prediction from the tumour BAF and correction of the LogR for GC-content biases^58^.

### Whole-exome sequencing

Genomic DNA was extracted from MEL006 PDX tumour samples (T0 and Tres) using the DNeasy Blood&Tissue kit (Qiagen). Exome sequencing was performed at the Genomics Core Facility, KU Leuven, Leuven, Belgium. Exome capture was performed with the SeqCap EZ Human Exome Library v3.0 (NimbleGen). These samples were subsequently sequenced in a paired end 151 bp run on a Novaseq instrument (Illumina, San Diego, CA) resulting in an average coverage of 120x.

### Exome sequencing analysis

Reads were aligned separately to the human (hs38) and mouse (mm10) reference genome using BWA mem (0.7.15-r1140, bwakit). Bamcmp (5cb0176) was used to deconvolve human and mouse reads^59^. Confident somatic short variants (PASS SNVs and indels) were subsequently called on the human BAM files using GATK (v4.1.2.0) following the Somatic Short Mutation calling Best Practice Workflow. Mutect2 was run in tumour-only mode on T0 and considering T0 as matched control for Tres. Variants were annotated using Funcotator. Variant allele frequencies were converted to cancer cell fractions in R using the estimated tumour purity and local total copy number^60^ as inferred by ASCAT on the corresponding SNP array data. Exposures of mutational signatures previously identified in whole-genome sequenced melanoma samples (COSMIC v3) were estimated using sigfit (v1.3.2) running 100 Markov Chain Monte Carlo (MCMC) simulations for 10,000 iterations including a 5,000 iteration burn-in^61^.

### Stem and lineage score assessment

Stemness and melanoma-lineage identities were inferred from scRNAseq data for different drug tolerant cell states^6^. For this purpose, the transcriptional activity of a melanoma stemness related gene set, which is induced upon exposure of melanoma cells to hESC medium^62^, was quantified for each single cell using AUCell based scoring^54^ and served as a proxy for stemness. Similarly, a gene set for melanoma-lineage identity^63^ was quantified in each single cell. Finally, drug tolerant cells of the undifferentiated, NCSC and hyperdifferentiated state were injected into a two-dimensional stemness/lineage space based on corresponding AUCell scores.

## GDNF ELISA

Cells were cultured and treated on a 96well plate in a total volume of 200ul. 100ul supernatant was used to determine GDNF concentration by ELISA according to manufacturer’s protocol (GDNF Emax® ImmunoAssay Systems, Promega). The plate with the remaining 100ul supernatant was used to determine cell numbers via Cell Titer Glo Assay according to manufacturer’s protocol (Promega).

### Sanger sequencing

Tumours were lysed and gDNA extracted with the NucleoSpin DNA RapidLyse kit according to manufacturer’s protocol (Machery-Nagel). NRAS exon3 was amplified out of 100ng gDNA via PCR and sent for Sanger sequencing. PCR protocol and primer sequences are available upon request.

### BRAF-splicing PCR

Between 200-300ng total RNA was used as input in a 20µl (total volume) SuperScript III (Invitrogen) RT reaction using oligo dT primers. We used 5µl of this cDNA reaction as input for the following PCR (50µl total volume): 5µl of Phusion High-Fidelity 10xPCR buffer (ThermoFisher), 1.5µl 10mM dNTP, 1.5µl 10 µM Fwd primer (BRAF trunc F-5’-GGCTCTCGGTTATAAGATGGC-3’), 1.5µl 10 µM Rev primer (BRAF trunc R-5’-ACAGGAAACGCACCATATCC-3’), 1µl 50mM MgSO4, 0.5µl Phusion High-Fidelity Taq polymerase and up to 45µl with water. Thermocycling conditions: 94°C 5min 1 cycle, 94°C 15sec_55°C 30sec_68°C 3min 38-40cycles, 68°C 5min 1cycle. Gel purify PCR-bands and Sanger-sequence with PCR primers^64^. The amplicon corresponding to BRAF WT is 2301bp.The alternative splicing variants are detected with amplicons of 1665bp for Δexon 4-8 and 1299bp for Δexon 2-8.

### Meta-analysis of resistance mechanisms to MAPK inhibition

DNA sequencing of BRAFi&MEKi resistant BRAF mutant melanoma patients of two centers was reassessed for the presence (genetic) or absence (non-genetic) of genetic alterations at resistance^17,23,28^. The percentage of genetic and non-genetic events was stratified by either number of samples or patients (Table S1, DNA summary).

### Meta-analysis of NCSC signature induction upon MAPK inhibition

Bulk-RNA sequencing data of matched treatment naïve and ON-treatment (BRAFi&MEKi) melanoma samples of three centers^17,23,28,65^ were interrogated for an induction of the NCSC gene expression program. Briefly, a gene expression average of the NCSC signature (n=42 genes) was calculated per sample. Then the ratios between ON-treatment over treatment-naïve NCSC averages were calculated for each sample/patient. An induction of 1.5-fold was considered to represent and NCSC induction (Table S1, RNA summary).

## ACKNOWLEDGMENTS

We thank Prof. G. Ghanem, LOCE-Institut J. Bordet, Université Libre de Bruxelles for providing us with the MM lines. We thank Göran Jönsson (Lund University) for the MM383 and WM852 cell lines. We thank Odessa Van Goethem and the excellent staff of the KULeuven PDX platform (TRACE) for their assistance with the PDX experiments. Flow cytometry/FACS was performed at the KU Leuven FACS Core Facility. Lukas Marcelis and Jasmin Garg provided histology support and help with mIHC, respectively. O.M-B. is supported by *12T1217N* project by FWO at the program under the Marie Skłodowska-Curie grant agreement no. 665501. P.K. received financial support from Marie Curie Individual Fellowship (H2020-MSCA-IF-2018) and J.F. from the Swedish Research Council. J.D. is supported by a postdoctoral fellowship from FWO. D.P. and N.V.D. received PhD fellowships from the VIB PhD international program and FWO-SB 1S79619N, respectively. This work was supported by FWO (#G.0929.16N), Stichting tegen kanker, Interreg (Skin-Huid), Melanoma Research Alliance (MRA, EIA#623591), KULeuven (C1 grant) to J-C.M., an OHSU Knight Cancer Center support grant from the National Institutes of Health (NIH P30-CA069533) to A.W.L., Infrastructure grants (type 1 funding from the Hercules Foundation – AKUL/13/41; Foundation Against Cancer project 2015-143) and grants from FWO (I001818N) and KU Leuven C14/18/092 to T.V. and enabled through access to the MRC eMedLab Medical Bioinformatics infrastructure, supported by the Medical Research Council (grant number MR/L016311/1).

## AUTHOR CONTRIBUTIONS

F.R., O.M-B. and A.R. designed and conducted experiments, acquired, analysed and interpreted the data. M.D., P.K., D.P. and N.V.D. conducted experiments and acquired data, M.D., and P.K. contributed to analysis and interpretation of the resulting data. E.L. provided help in setting up and interpret the PDX experiments. J.P., J.D. D.L. and T.V. helped generate, analysed and interpreted the shallow WGS, WES and SNP array data. G.B. conducted SMARTseq2 and qPCR experiments. O.B., F.B., M.L., H.R. and J.vd.O. provided human samples and pathology support. I.vd.B. and S.vd.B. conducted DNA FISH BRAFamp analysis and TruSight data interpretation, respectively. J.F. conducted mIHC and A.W.L. interpreted the data. All authors read and edited the manuscript. F.R. and J.–C.M. conceptualized, designed research studies and wrote the manuscript.

