## Supplemental Data for "A neural crest stem cell-like state drives nongenetic resistance to targeted therapy in melanoma"

### Figure S1

## A

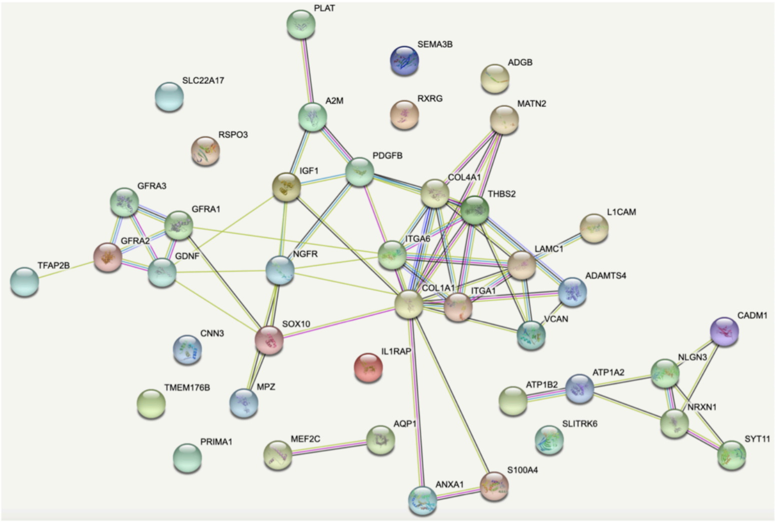


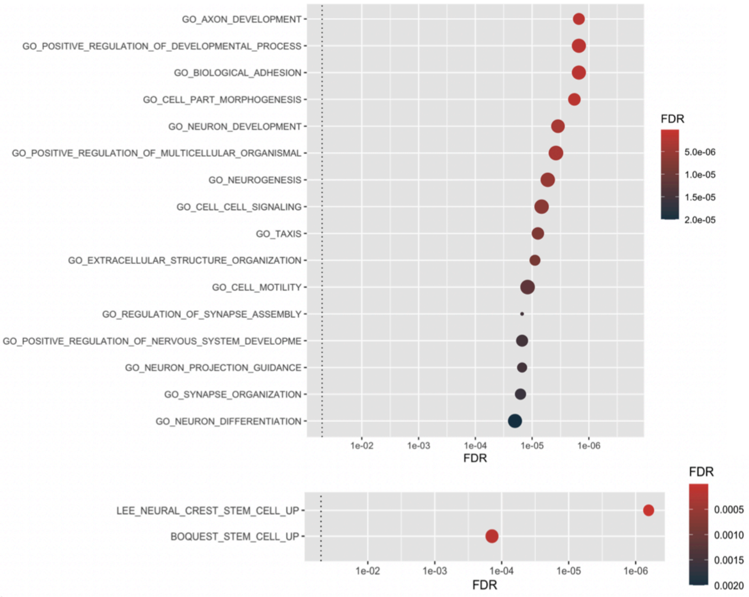


**Reference publications (PubMed)**

| publication | (year) title | count in network | strength | false discovery rate |
|---|---|---|---|---|
| PMID:20116071 | (2010) Genetic association of the GDNF alpha-receptor gen... | 4 of 4 | 2.67 | 5.81e-05 |
| PMID:10974669 | (2000) GDNF family receptors in the embryonic and postnat... | 3 of 3 | 2.67 | 0.00044 |
| PMID:9843911 | (1998) Regulation of c-ret expression by retinoic acid in rat ... | 2 of 2 | 2.67 | 0.0059 |
| PMID:9808338 | (1998) Glial cell line-derived neurotrophic factor (GDNF): a ... | 2 of 2 | 2.67 | 0.0059 |
| PMID:9790508 | (1998) COL1A1-PDGFB fusion in a ring chromosome 4 foun... | 2 of 2 | 2.67 | 0.0059 |

**Biological Process (Gene Ontology)**

| GO-term | description | count in network | strength | false discovery rate |
|---|---|---|---|---|
| GO:0099174 | regulation of presynapse organization | 2 of 6 | 2.19 | 0.0022 |
| GO:0035860 | glial cell-derived neurotrophic factor receptor signaling path... | 2 of 8 | 2.07 | 0.0032 |
| GO:0097105 | presynaptic membrane assembly | 2 of 9 | 2.02 | 0.0038 |
| GO:0097104 | postsynaptic membrane assembly | 2 of 9 | 2.02 | 0.0038 |
| GO:0038063 | collagen-activated tyrosine kinase receptor signaling pathw... | 2 of 9 | 2.02 | 0.0038 |

## B

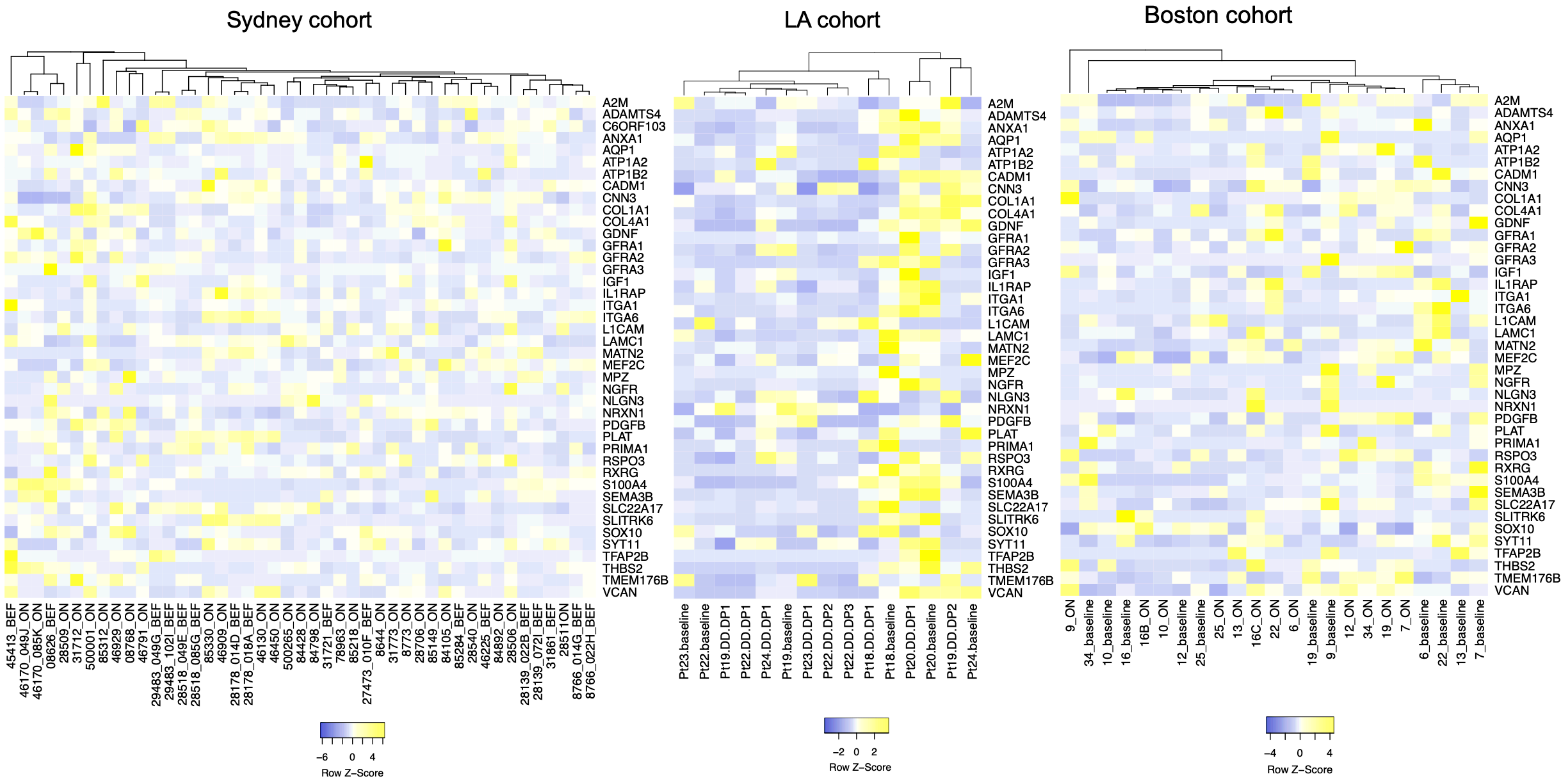

**Figure S1 (linked to Figure 1)**

A) Protein network analysis and enriched terms (string-db) as well as significant gene enrichment terms (hypeR) showing the nature of the NCSC gene signature (n=42)

B) Heatmaps depict gene expression (bulk RNA-seq) of the NCSC signature in three BRAFi+MEKi treated patient cohorts and matching treatment naïve samples (BEF=BEFORE, baseline). Data are from three different clinical centers: Melanoma Institute Australia (Sydney, Australia), UCLA Dermatology (Los Angeles, USA) and Massachusetts General Hospital (Boston, USA)

### Figure S2

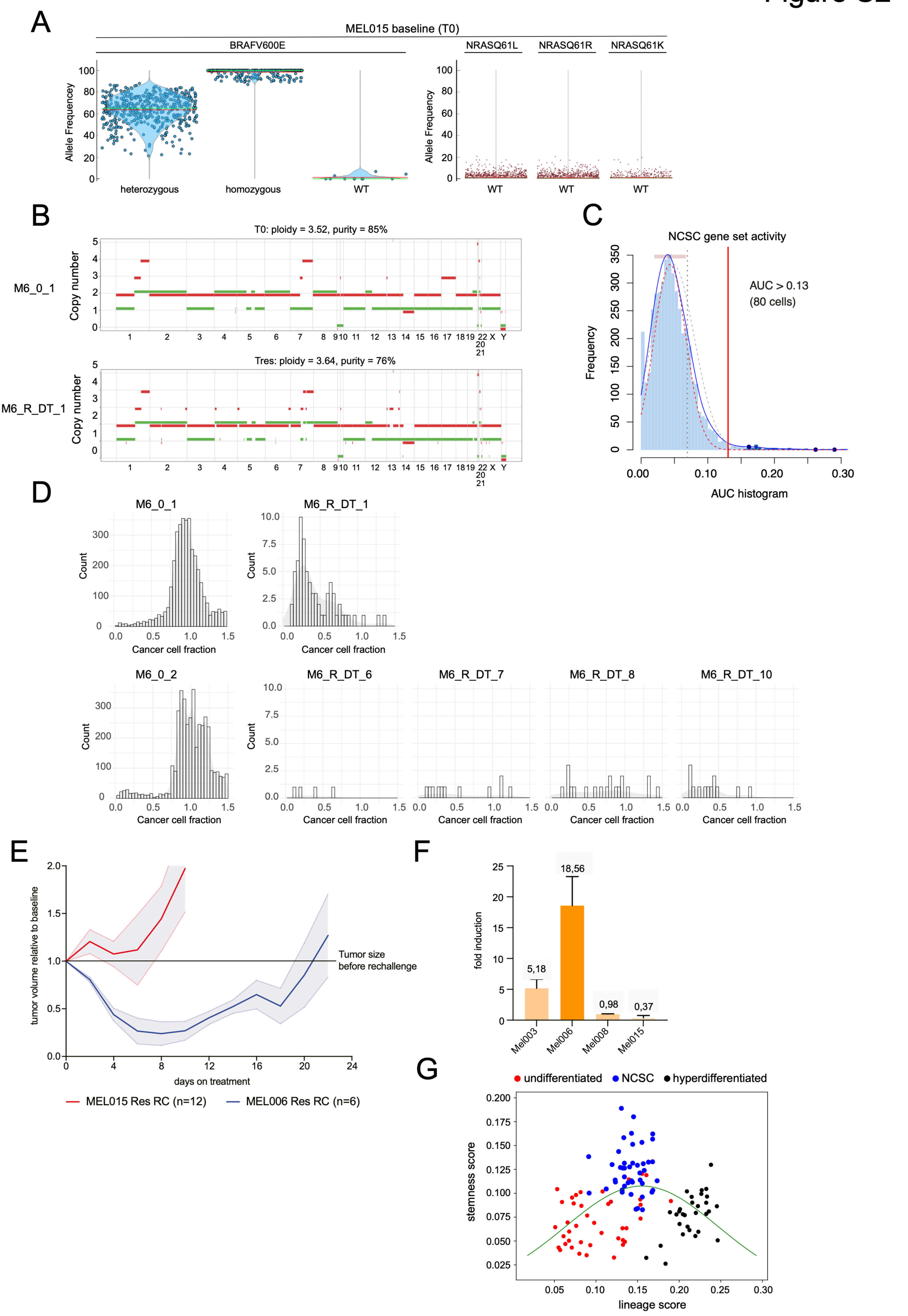
A
MEL015 baseline (T0)
BRAFV600E
NRASQ61L
NRASQ61R
NRASQ61K
Allele Frequencey
Allele Frequency
heterozygous
homozygous
WT
B
T0: ploidy = 3.52, purity = 85%
M6_0_1
Copy number
Tres: ploidy = 3.64, purity = 76%
M6_R_DT_1
C
NCSC gene set activity
Frequency
AUC > 0.13
(80 cells)
AUC histogram
D
M6_0_1
M6_R_DT_1
M6_0_2
M6_R_DT_6
M6_R_DT_7
M6_R_DT_8
M6_R_DT_10
Count
Cancer cell fraction
E
tumor volume relative to baseline
Tumor size
before rechallenge
days on treatment
MEL015 Res RC (n=12)
MEL006 Res RC (n=6)
F
fold induction
5,18
18,56
0,98
0,37
Mel003
Mel006
Mel008
Mel015
G
undifferentiated
NCSC
hyperdifferentiated
stemness score
lineage score

**Figure S2 (linked to Figure 2)**

A) Single-cell targeted DNA sequencing (Tapestri, missionbio) of a MEL015 treatment naïve lesion (about 3.5k single cells). Allele frequencies for BRAFV600E and NRASQ61K/L/R are shown.

B) Allele-specific copy number plots of T0 and Tres (MEL006). Major and minor alleles are indicated in red and green, respectively.

C) Frequency plot shows the distribution of NCSC AUCell scores for MEL006 and MEL015 single cells and the applied cut-off: AUCell score > 0.13, which renders 80 NCSC-state cells.

D) Histogram of cancer cell fractions for *de novo* TRes SNVs inferred from n=5 resistant tumour samples and their corresponded drug naïve counterpart. An estimated cancer cell fraction of .45 suggests the variant is present in 45% of tumour cells.

E) Tumour-volume in function of time of therapy resistant MEL06 (blue, n=6) and MEL015 (red, n=12) PDX lesions exposed to BRAF/MEKi after a drug-holiday period. Data is represented by mean (thick line) ±SEM (filled area).

F) The percentage of NGFR/AQP1-double-positive cells (NCSCs) was determined by immunohistochemistry at MRD and before treatment (T0) in PDX lesions from group 2: MEL03, MEL06, MEL08 and MEL015. Data represent the mean fold induction in MRD compared to T0 ±SD.

G) Stemness and melanoma-lineage identities were inferred from scRNAseq data for different drug tolerant cell states. Drug tolerant cells of the undifferentiated, NCSC and hyperdifferentiated state were injected into a two-dimensional stemness/lineage space based on corresponding AUCell scores. The green line represents the Gaussian data fit.

### Figure S3

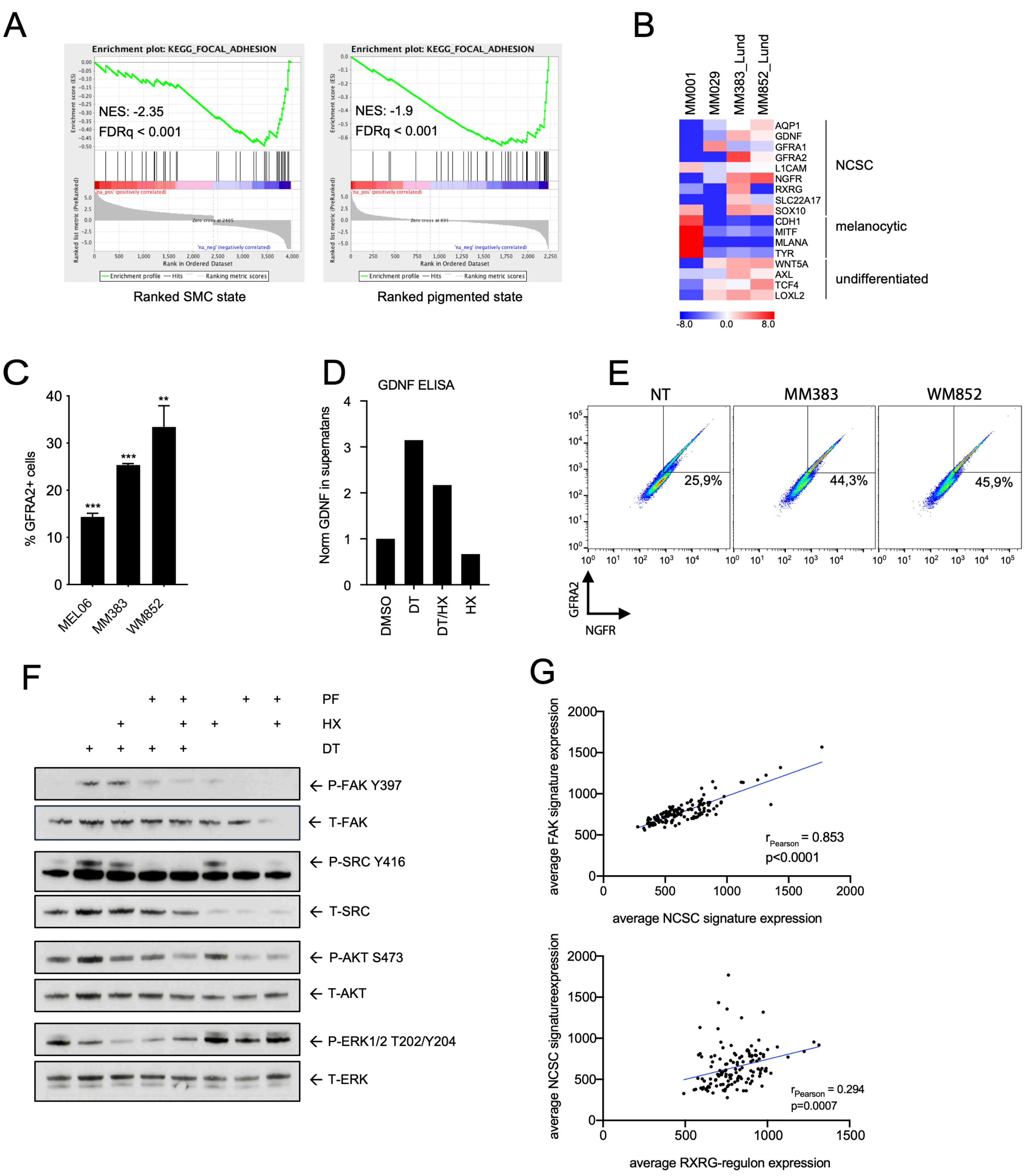

A
Enrichment plot: KEGG_FOCAL_ADHESION
NES: -2.35
FDRq < 0.001
Ranked SMC state
Enrichment plot: KEGG_FOCAL_ADHESION
NES: -1.9
FDRq < 0.001
Ranked pigmented state
B
MM001
MM029
MM383_Lund
MM852_Lund
AQP1
GDNF
GFRA1
GFRA2
L1CAM
NGFR
RXRG
SLC22A17
SOX10
CDH1
MITF
MLANA
TYR
WNT5A
AXL
TCF4
LOXL2
NCSC
melanocytic
undifferentiated
-8.0
0.0
8.0
C
% GFRA2+ cells
MEL06
MM383
WM852
D
GDNF ELISA
Norm GDNF in supernatans
DMSO
DT
DT/HX
HX
E
NT
MM383
WM852
25,9%
44,3%
45,9%
GFRA2
NGFR
F
PF
HX
DT
P-FAK Y397
T-FAK
P-SRC Y416
T-SRC
P-AKT S473
T-AKT
P-ERK1/2 T202/Y204
T-ERK
G
average FAK signature expression
average NCSC signature expression
$r_{Pearson}$ = 0.853
p<0.0001
average NCSC signatureexpression
average RXRG-regulon expression
$r_{Pearson}$ = 0.294
p=0.0007

**Figure S3 (linked to Figure 3)**

A) Gene set enrichment analysis (GSEA) shows absence of enrichment of Focal Adhesion signalling (KEGG pathway) in the SMC and pigmented DTC states[6]. NES, normalized enrichment score; FDR, false discovery rate.

B) Heat map for key marker genes of the Neural Crest Stem Cell (NCSC), melanocytic and undifferentiated cell populations, comparing melanoma lines MM001 (melanocytic), MM029 (undifferentiated), MM383 and WM852_lund (NCSC).

C) Quantitative flow cytometry analysis of GFRA2+ cells in MEL006, MM383 and WM852 cell cultures.

D) GDNF ELISA measuring GDNF secretion in the cellular supernatant of MM383 cells treated with DT (100nM/20nM) and the RXR antagonist HX531 (2uM) for 72h. The measured GDNF concentrations were corrected for cell number and normalized to DMSO control.

E) FACS plots showing the gating strategy of Figure 3H.

F) Western blot analysis of levels of pan- and phosphorylated-FAK, AKT and ERK in MM099 treated for 72h with DT (100nM/20nM), RXR antagonist HX531 (2uM) and FAK-inhibitor PF562271 (1uM). SRC phosphorylation was used as marker for FAK signalling activity.

G) Scatter plots showing significant correlations between the FAK and NCSC gene expression signatures in bulk RNAseq data from melanoma lesions exposed to MAPK-inhibitors (cohort from the Melanoma Institute Australia, Sydney). A significant correlation was also observed between the NCSC gene expression signature and RXRG-regulon activity.

### Figure S4

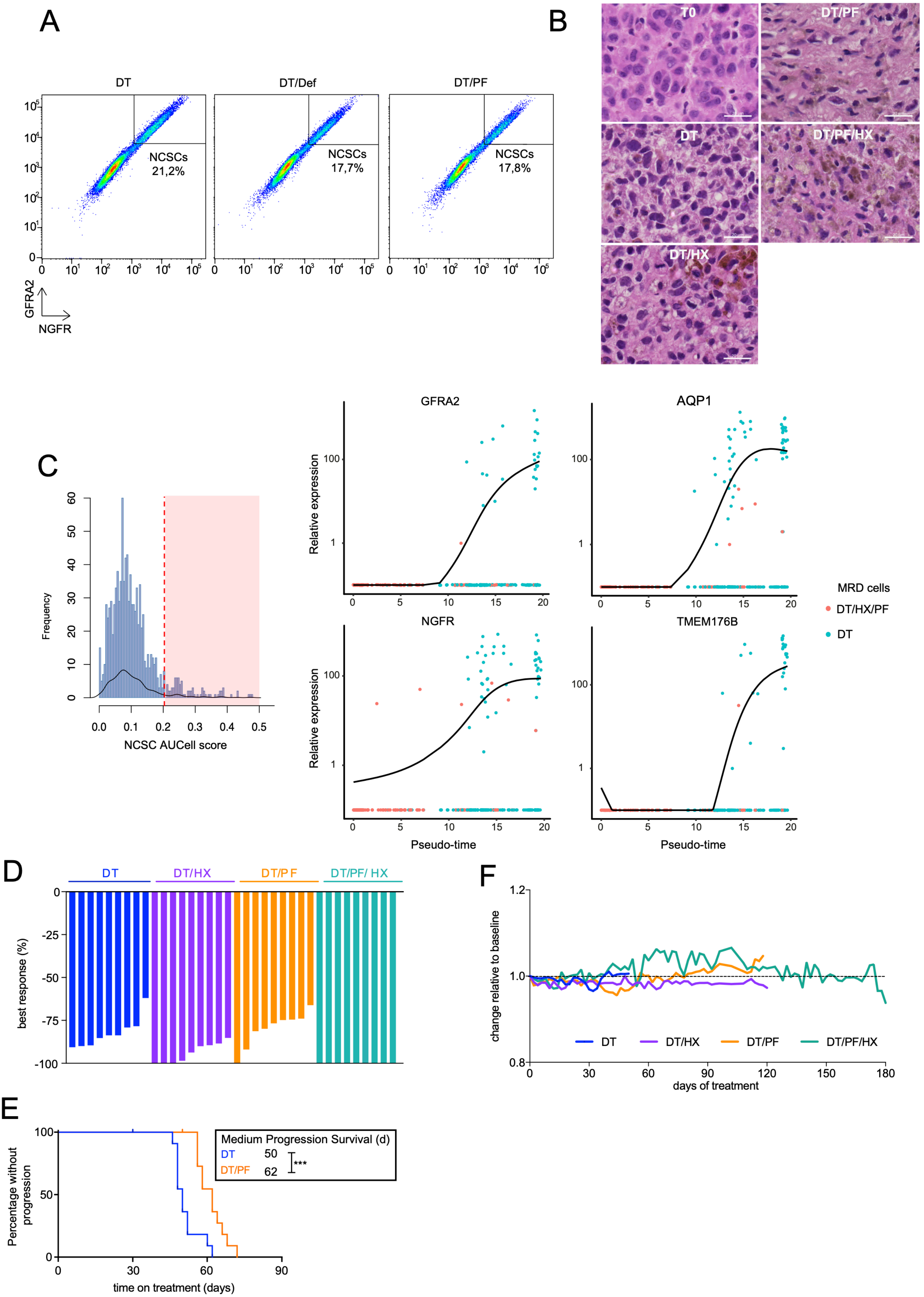
A
B
DT
DT/Def
DT/PF
NCSCs 21,2%
NCSCs 17,7%
NCSCs 17,8%
GFRA2
NGFR
T0
DT/PF
DT
DT/PF/HX
DT/HX
C
Frequency
NCSC AUCell score
GFRA2
AQP1
NGFR
TMEM176B
Relative expression
Pseudo-time
MRD cells
DT/HX/PF
DT
D
DT
DT/HX
DT/PF
DT/PF/ HX
best response (%)
F
change relative to baseline
DT
DT/HX
DT/PF
DT/PF/HX
days of treatment
E
Percentage without progression
Medium Progression Survival (d)
DT 50
DT/PF 62
***
time on treatment (days)

**Figure S4 (linked to Figure 4)**

A) Flow cytometry plots showing gating strategy for main Figure 4A.

B) Hematoxilin and eosin staining (H&E) before treatment (T0) and ON treatment with different drug combinations; dabrafenib-trametinib alone (DT), DT plus HX531 (DT/HX), DT plus FAK inhibitor, PF562271 (DT/PF) and the quadruple combination (DT/PF/HX) at MRD. Scale bar=50 μm (above) and bar=20 μm (below).

C) AUCell score frequency distribution plot for NCSC activity in single-cell RNA sequencing data from MRD lesions. Inclusion criteria (red surface) AUCell score > 0.2 (Figure 4E). NCSC marker expression examples (GFRA2, AQP1, NGFR, TMEM176B) during MRD inferred by pseudotime ordering (monocle pipeline).

D) Waterfall plot showing best response (%) of MEL006 PDX model treated with; dabrafenib-trametinib (DT) alone (n = 9), DT plus HX531 (DT/HX), DT plus PF562271 (DT/PF) and the quadruple combination (DT/PF/HX) at day 14 after treatment. In the DT/PF/HX-treated group all the mice achieved a complete response.

E) Kaplan-Meier curve for MEL015 mice treated with dabrafenib-trametinib alone (DT, n = 10) and DT plus FAK inhibitor, PF562271 (DT/PF, n = 10). Median time to progression was 50 days for DT and 62 days for DT/PF. Log rank (Mantel-Cox) for DT versus DT/PF: $p < 0.001$ (***).

F) Body weight curves relative to baseline. MEL006 mice treated with dabrafenib-trametinib (DT), DT plus HX531 (DT/HX), DT plus PF562271 (DT/PF) and the quadruple combination (DT/PF/HX). The drugs were administered daily by oral gavage.

Figure S5

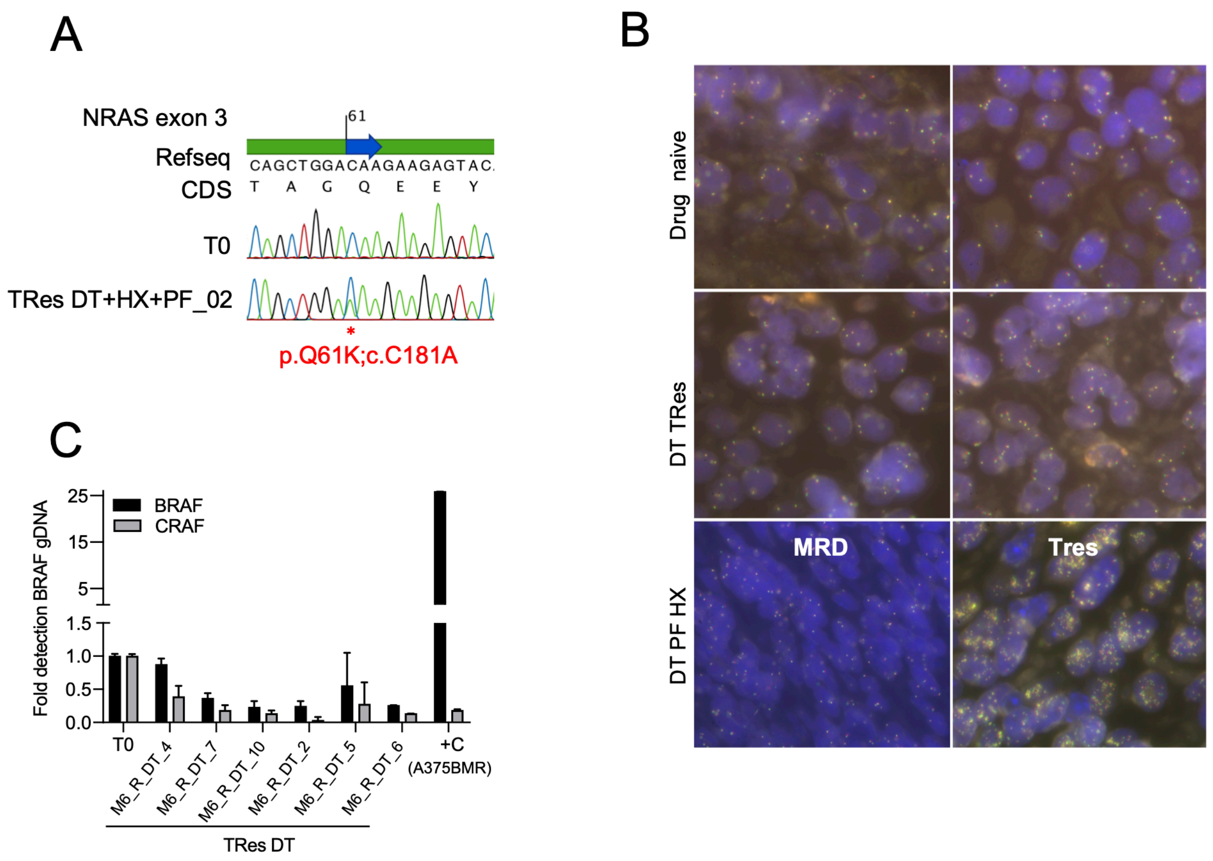

A
NRAS exon 3
61
Refseq
CAGCTGGACAAGAAGAGTAC
CDS
T A G Q E E Y
T0
TRes DT+HX+PF_02
*
p.Q61K;c.C181A
B
Drug naive
DT TRes
DT PF HX
MRD
Tres
C
Fold detection BRAF gDNA
BRAF
CRAF
25
15
5
1.5
1.0
0.5
0.0
T0
M6_R_DT_4
M6_R_DT_7
M6_R_DT_10
M6_R_DT_2
M6_R_DT_5
M6_R_DT_6
+C
(A375BMR)
TRes DT

**Figure S5 (linked to Figure 5)**

A) Sanger sequencing of NRAS exon3 genomic DNA in an untreated MEL006 PDX model (T0) and a tumour resistant to DT/PF/HX. The image shows the sequencing data around codon 61.

B) Representative pictures of DNA FISH analysis for BRAF SPEC BRAF Dual Colour Break Apart Probe (ZytoVision) of PDX tissue sections before treatment (T0, left) and tumours resistant to DT/PF/HX treatment (Tres DT/HX/PF, right).

C) Quantitative PCR analysis of *BRAF* and *CRAF* copy number was performed using genomic DNA from DT-resistant lesions (MEL006). (RQ=relative quantity of DNA copies compared to T0).

**Table S1 (linked to Figure 1)**

Sample and patient details of genetic analyses addressing resistance mechanism.

| study | treatment | Patient | sample ID | Res_mechanism_call | alteration | (Best) Response or RECIST |
|---|---|---|---|---|---|---|
| Sydney | BM | NC_1 | 8766_022H PreC | genetic | MEK1 G128D, MEK2 C125S | SD |
| Sydney | BM | NC_1 | 8773 PostC | genetic | MEK1 G128D, MEK2 C125S | SD |
| Sydney | BM | NC_1 | 28509 ProgC | genetic | MEK1 G128D, MEK2 C125S | SD |
| Sydney | BM | NC_1 | 08766 PreC_014G | genetic | MEK1 G128D, MEK2 C125S | SD |
| Sydney | BM | NC_1 | 28506 ProgC | genetic | MEK1 G128D, MEK2 C125S | SD |
| Sydney | BM | NC_1 | 28511 ProgC | genetic | MEK1 G128D, MEK2 C125S | SD |
| Sydney | BM | NC_10 | 46450 ProgC | genetic | BRAF amp | SD |
| Sydney | BM | NC_10 | 46130 ProgC | genetic | BRAF amp | SD |
| Sydney | BM | NC_10 | 46909 Progc | genetic | BRAF amp | SD |
| Sydney | BM | NC_10 | 84892 ProgC | genetic | BRAF amp | SD |
| Sydney | BM | NC_10 | 500001 ProgC | genetic | BRAF amp | SD |
| Sydney | BM | NC_10 | 46929 ProgC | genetic | BRAF amp | SD |
| Sydney | BM | NC_10 | 85330 ProgC | genetic | BRAF amp | SD |
| Sydney | BM | NC_2 | 31721 PreC | genetic | NRAS Q61K | SD |
| Sydney | BM | NC_2 | 31773 PostC | genetic | NRAS Q61K | SD |
| Sydney | BM | NC_2 | 85218 ProgC | genetic | NRAS Q61K | SD |
| Sydney | BM | NC_2 | 85149 ProgC | genetic | NRAS Q61K | SD |
| Sydney | BM | NC_3 | 28518_ 085G PreC | genetic | MEK2 C125S | SD |
| Sydney | BM | NC_3 | 28518_049E PreC | genetic | MEK2 C125S | SD |
| Sydney | BM | NC_3 | 28706 ProgC | genetic | MEK2 C125S | SD |
| Sydney | BM | NC_4 | 08626 PreC | non-genetic | nil | PR |
| Sydney | BM | NC_4 | 46791 ProgC | non-genetic | nil | PR |
| Sydney | BM | NC_5 | 46225 PreC | genetic | MEK2 E207K | PR |
| Sydney | BM | NC_5 | 78963 ProgC | genetic | MEK2 E207K | PR |
| Sydney | BM | NC_6 | 85284 PreC | genetic | NRAS Q61K | SD |
| Sydney | BM | NC_6 | 85312 PostC | genetic | NRAS Q61K | SD |
| Sydney | BM | NC_6 | 84105 ProgC | genetic | NRAS Q61K | SD |
| Sydney | BM | NC_7 | 46170_085K ProgC off | genetic | BRAF amp | PR |
| Sydney | BM | NC_7 | 45413 PreC | genetic | BRAF amp | PR |
| Sydney | BM | NC_7 | 46170_049J ProgC (Off drug) | genetic | BRAF amp | PR |
| Sydney | BM | NC_9 | 29483_049G PreC | genetic | BRAF amp | PR |
| Sydney | BM | NC_9 | 84428 ProgC | genetic | BRAF amp | PR |
| Sydney | BM | NC_9 | 29483_102I PreC | genetic | BRAF amp | PR |
| Sydney | BM | NC_9 | 500265 ProgC | genetic | BRAF amp | PR |
| Sydney | BM | NC_9 | 84798 ProgC | genetic | BRAF amp | PR |
| LA | BM | Pt18 | Pt18-DD-DP1 | genetic | NRAS SNV/INDEL and PTEN SNV/INDEL | -70% |
| LA | BM | Pt19 | Pt19-DD-DP1 | genetic | KRAS SNV/INDEL and PTEN loss | -51% |
| LA | BM | Pt19 | Pt19-DD-DP2 | genetic | KRAS SNV/INDEL | -51% |
| LA | BM | Pt20 | Pt20-DD-DP1 | non-genetic | nil | -60% |
| LA | BM | Pt21 | Pt21-DD-DP1 | genetic | NRAS mut.allele CN gain | -40% |
| LA | BM | Pt21 | Pt21-DD-DP2 | genetic | NRAS mut.allele CN gain | -40% |
| LA | BM | Pt22 | Pt22-DD-DP1 | genetic | BRAF amp. and DUSP4 loss | NA |
| LA | BM | Pt22 | Pt22-DD-DP2 | genetic | BRAF amp. and DUSP4 loss | NA |
| LA | BM | Pt22 | Pt22-DD-DP3 | genetic | BRAF amp. and DUSP4 loss | NA |
| LA | BM | Pt23 | Pt23-DD-DP1 | non-genetic | nil | -30% |
| LA | BM | Pt23 | Pt23-DD-DP2 | non-genetic | nil | -30% |
| LA | BM | Pt23 | Pt23-DD-DP3 | non-genetic | nil | -30% |
| LA | BM | Pt23 | Pt23-DD-DP4 | non-genetic | nil | -30% |
| LA | BM | Pt23 | Pt23-DD-DP5 | non-genetic | nil | -30% |
| LA | BM | Pt23 | Pt23-DD-DP6 | non-genetic | nil | -30% |
| LA | BM | Pt24 | Pt24-DD-DP1 | non-genetic | nil | -100% |
| LA | BM | Pt28 | Pt28-DD-DP3 | genetic | BRAF amp. and CDKN2A loss | 28% |
| LA | BM | Pt28 | Pt28-DD-DP9 | genetic | BRAF amp. and CDKN2A loss | 28% |
| LA | BM | Pt28 | Pt28-DD-DP10 | genetic | BRAF amp. and CDKN2A loss | 28% |
| LA | BM | Pt28 | Pt28-DD-DP4 | genetic | CDKN2A loss | 28% |
| LA | BM | Pt28 | Pt28-DD-DP1 | genetic | CDKN2A loss | 28% |
| LA | BM | Pt28 | Pt28-DD-DP5 | genetic | CDKN2A loss | 28% |
| LA | BM | Pt28 | Pt28-DD-DP2 | non-genetic | nil | 28% |
| LA | BM | Pt29 | Pt29-DD-DP1 | genetic | CDKN2A loss and PTEN loss | -9% |

**treatment**
**BM=BRAFi+MEKi**

| patient_ID | sample comparison (ON vs BEFORE) | ratio NCSS | treament | City |
|---|---|---|---|---|
| 16 | 16C | 3,088 | BM | Boston |
| 12 | 12B | 2,474 | BM | Boston |
| 9 | 9B | 2,014 | BM | Boston |
| 16 | 16C | 1,699 | BM | Boston |
| 19 | 19B | 1,028 | BM | Boston |
| 10 | 10B | 0,968 | BM | Boston |
| 13 | 13B | 0,868 | BM | Boston |
| 7 | 7B | 0,809 | BM | Boston |
| 25 | 25C | 0,800 | BM | Boston |
| 34 | 34B | 0,627 | BM | Boston |
| 22 | 22C | 0,576 | BM | Boston |
| 6 | 6B | 0,433 | BM | Boston |
| Pt19 | Pt19-DD-DP2 | 4,426 | BM | LA |
| Pt22 | Pt22-DD-DP3 | 2,013 | BM | LA |
| Pt22 | Pt22-DD-DP2 | 1,946 | BM | LA |
| Pt20 | Pt20-DD-DP1 | 1,087 | BM | LA |
| Pt22 | Pt22-DD-DP1 | 1,048 | BM | LA |
| Pt18 | Pt18-DD-DP1 | 0,876 | BM | LA |
| Pt23 | Pt23-DD-DP1 | 0,606 | BM | LA |
| Pt19 | Pt19-DD-DP1 | 0,468 | BM | LA |
| Pt24 | Pt24-DD-DP1 | 0,316 | BM | LA |
| NC_2 | SMU-030 | 1,858 | BM | Sydney |
| NC_1 | WMD-022 | 1,637 | BM | Sydney |
| NC_10 | WMD-017 | 1,543 | BM | Sydney |
| NC_6 | SMU-034 | 1,509 | BM | Sydney |
| NC_2 | SMU-030 | 1,467 | BM | Sydney |
| NA_1 | SMU-029 | 1,444 | BM | Sydney |
| NC_6 | SMU-034 | 1,270 | BM | Sydney |
| NC_1 | WMD-022 | 1,256 | BM | Sydney |
| NC_3 | MTP-034 | 1,140 | BM | Sydney |
| NC_10 | WMD-017 | 1,047 | BM | Sydney |
| NC_2 | SMU-030 | 1,042 | BM | Sydney |
| NC_1 | WMD-022 | 1,035 | BM | Sydney |
| NC_1 | WMD-022 | 1,011 | BM | Sydney |
| NC_10 | WMD-017 | 1,009 | BM | Sydney |
| NC_10 | WMD-017 | 0,924 | BM | Sydney |
| NC_10 | WMD-017 | 0,913 | BM | Sydney |
| NC_10 | WMD-017 | 0,902 | BM | Sydney |
| NC_9 | SMU-028 | 0,862 | BM | Sydney |
| WMD-039 | WMD-039 | 0,846 | BM | Sydney |
| NC_4 | WMD-033 | 0,841 | BM | Sydney |
| CCR_24 | WMD-013 | 0,821 | BM | Sydney |
| NC_7 | MTP-073 | 0,764 | BM | Sydney |
| NC_7 | MTP-073 | 0,735 | BM | Sydney |
| CCR_24 | WMD-013 | 0,728 | BM | Sydney |
| NC_9 | SMU-028 | 0,692 | BM | Sydney |
| NC_10 | WMD-017 | 0,648 | BM | Sydney |
| NC_5 | SMU-020 | 0,585 | BM | Sydney |
| NC_9 | SMU-028 | 0,506 | BM | Sydney |

**NCSC signature**

A2M
ADAMTS4
C6ORF103
ANXA1
AQP1
ATP1A2
ATP1B2
CADM1
CNN3
COL1A1
COL4A1
GDNF
GFRA1
GFRA2
GFRA3
IGF1
IL1RAP
ITGA1
ITGA6
L1CAM
LAMC1
MATN2
MEF2C
MPZ
NGFR
NLGN3
NRXN1
PDGFB
PLAT
PRIMA1
RSPO3
RXRG
S100A4
SEMA3B
SLC22A17
SLITRK6
SOX10
SYT11
TFAP2B
THBS2
TMEM176B
VCAN
